# Identification of clofibric acid as a SYVN1 ligand for PROTAC development

**DOI:** 10.64898/2026.01.26.701774

**Authors:** Julia Warren, Anandarao Munakala, Keith D. Zientek, Kilsun Kim, Phil A. Wilmarth, Ashok P. Reddy, Bingbing X. Li, Xiangshu Xiao

## Abstract

Targeted protein degradation (TPD) is an emerging therapeutic modality for numerous diseases. PROteolysis-TArgeting Chimeras (PROTACs) represent a potentially generalizable strategy to achieve TPD. A PROTAC is composed of a ligand for a protein of interest, a linker and a ligand for E3 ligase. As such, PROTACs can bring the E3 ligase into the close proximity of a protein target leading to polyubiquitination followed by target protein degradation. While the human genome encodes over 600 E3 ligases, only a handful of them have been harnessed for developing PROTACs. In order to expand the repertoire of E3 ligases for PROTAC development, we developed clickable photoaffinity probes based on clinically used drugs and metabolites to identify potential E3 ligases as the targets. In this paper, we report the discovery of clofibric acid with a molecular weight of only 214 Daltons as a ligand for synoviolin (SYVN1). We demonstrate its utility by developing clofibric acid-based BRD4 PROTACs. The linker length and architecture play a critical role in target degradation efficiency. The clofibric acid-derived BRD4 PROTACs achieve selective BRD4 degradation in a SYVN1-dependent manner. Our findings establish clofibric acid as a robust addition to the TPD toolbox, offering a novel E3 ligase recruitment strategy for the development of next-generation degraders.

**TOC Graphics:** 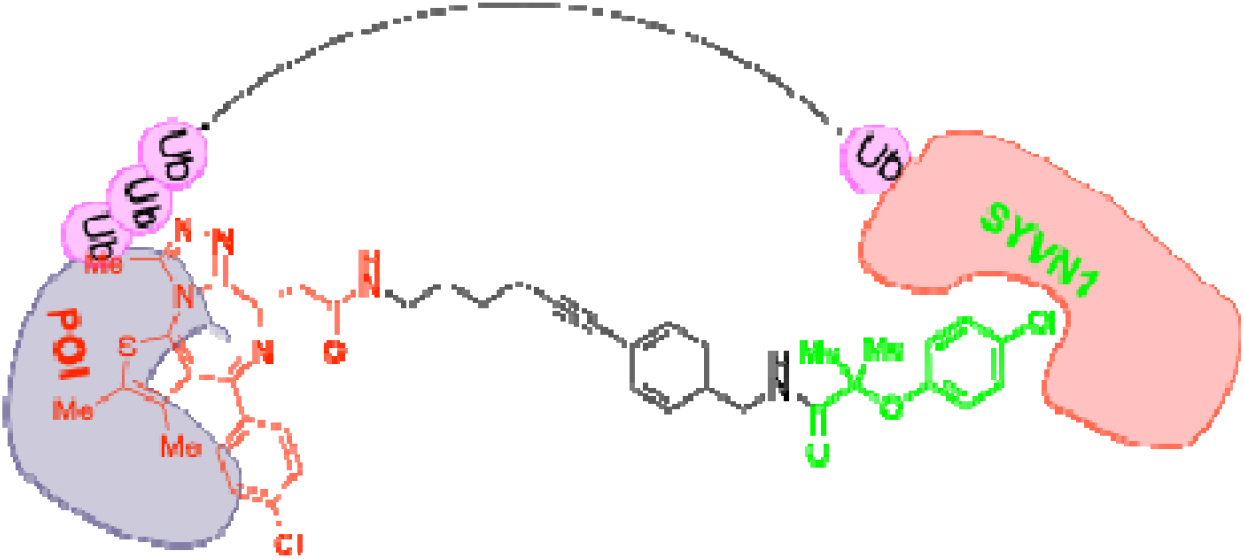

## INTRODUCTION

Targeted protein degradation (TPD) is an emerging therapeutic modality by degrading a target protein in its entirety.^1^ PROteolysis-TArgeting Chimera (PROTAC) is the most commonly used strategy to achieve TPD. A PROTAC is a heterobifunctional molecule containing a ligand for a protein of interest (POI), a ligand for a E3 ligase and a linker to connect these two ligands together.^2^ As such, a PROTAC can induce formation of a functional ternary complex containing the POI, a E3 ligase and PROTAC itself.^3^ Upon formation of the ternary complex, the target POI becomes polyubiquitinated by the recruited E3 ligase (Figure 1A), which is then followed by proteolysis by the ubiquitin-proteasome system (UPS).^2^ After degradation, the liberated PROTAC molecule can initiate future rounds of protein degradation, rendering such mechanism of degradation catalytic with event-driven pharmacology rather than occupancy-driven pharmacology.^2^ Furthermore, since the protein-targeting ligand in a PROTAC only requires binding with the target POI, it has tremendous potential to drug many undruggable proteins including scaffold proteins and transcription factors, where *bona fide* orthosteric sites do not exist or are undruggable.^2, 4^ These unique properties have fueled great interest in the community to pursue PROTACs as novel therapeutics and indeed over 40 different PROTACs have entered various stages of clinical trials.^5^

**Figure 1.**
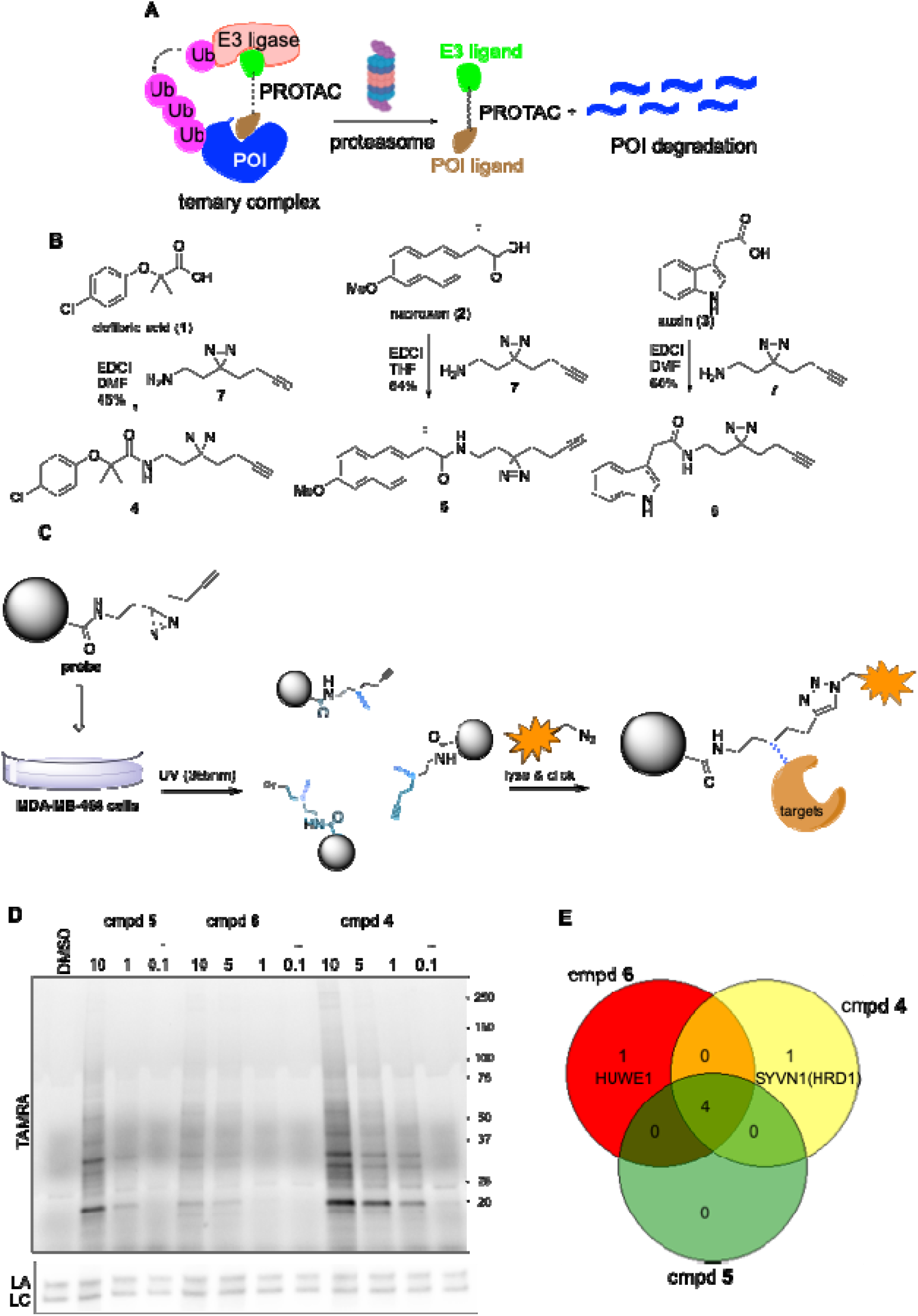
Identification of SYVN1 as a potential target of clofibric acid. (A) A schematic diagram showing the mechanism of action of PROTACs. (B) Synthesis of probes **4-6** from their corresponding parent carboxylic acids **1-3**. (C) A schematic flow diagram showing the target identification using clickable photoaffinity probes from live MDA-MB-468 cells. (D) Probes **4-6** labeled unique protein targets in MDA-MB-468 cells. The cells were treated with different concentrations of probes **4-6** for 30 min. Then the cells were UV-irradiated at 365 nm for 5 min. The cells were collected to prepare the lysates, which were then clicked with TAMRA-N_3_. The clicked lysates were separated on an SDS-PAGE for in-gel fluorescence analysis. The proteins were further transferred to a nitrocellulose membrane for western blotting with anti-lamin A (LA) as a loading control. (E) Identification of E3 ligases as potential targets of probes **4-6** by LC-MS/MS analysis. The identified E3 ligases from different probes are presented as a Venn diagram.

The human genome encodes more than 600 E3 ligases.^6^ However, only a handful of E3 ligases (∼3%) have been successfully hijacked for PROTAC development.^7-9^ Examples include CRBN (cereblon),^10^ VHL (Von Hippel-Lindau),^11^ IAPs (inhibitor of apoptosis),^12^ DCAF1 (DDB1 and CUL4 associated factor 1),^13^ DCAF11 (DDB1 and CUL4 associated factor 11),^14^ DCAF16 (DDB1 and CUL4 associated factor 16),^15^ RNF114 (ring finger protein 14),^16^ MDM2 (mouse double minute 2),^17^ FEM1B (Fem-1 homolog B),^18^ and FBXO22 (F-box only protein 22).^19^ This leaves the vast majority of the E3 ligases (97%) remaining to be hijacked for TPD. The two workhorse E3 ligases are CRBN and VHL. Identification of additional E3 ligases for TPD has at least following three advantages: 1) Expanding the druggable proteome by pairing a right target with a right E3 ligase. Not all E3 ligases are created equally. Some E3 ligases, when conjugated to a target, could not induce target degradation.^20^ 2) Achieving tissue selectivity by hijacking right E3 ligases in the right tissue to be targeted. Bcl-xL targeting PROTACs were developed by conjugating a Bcl-xL inhibitor with a VHL ligand to avoid toxicity associated with Bcl-xL inhibitors in platelets due to limited VHL expression in platelets.^21^ 3) Achieving subcellular selectivity to degrade the target only in the intended subcellular compartment where E3 ligase is located. DCAF16 was shown to induce target degradation only in the nucleus without affecting the target localized in the cytosol.^15^ With these potential advantages, expanding the repertoire of druggable E3 ligases will ultimately increase the safety of PROTAC molecules for clinical use. In this paper, we describe the identification of clofibric acid (**1**, Figure 1B) as a small molecule ligand for E3 ligase synoviolin 1 (SYVN1, aka HRD1). We further demonstrate its utility for developing PROTACs to achieve TPD.

## RESULTS AND DISCUSSION

### Identification of SYVN1 as a target of clofibric acid

To identify novel ligands for E3 ligases for PROTAC development, it is highly desirable to have the ligands that are small in molecular weight given that the final PROTAC molecules contains a POI ligand and an additional linker. Indeed, due to the smaller size of CRBN ligands, the vast majority of PROTACs in clinical development are based on CRBN ligands.^22^ We hypothesized that small fragment-like molecules from clinically used drugs and endogenous metabolites may possess hitherto unknown E3 ligase-binding activities. The advantages of such small molecules include known safety profiles in humans and potential oral bioavailability. To test this hypothesis, we began with clofibric acid (**1**), naproxen (**2**) and auxin (**3**) (Figure 1B). All these 3 compounds are carboxylic acids connected to a hydrophobic aromatic core through a short linker. Clofibric acid is the active metabolite of cholesterol-lowering drug clofibrate that is known to target peroxisome proliferator-activated receptor α (PPARα).^23, 24^ Naproxen is a non-steroid anti-inflammatory drug (NSAID) targeting cyclooxygenase 2 (COX-2)^25^ while auxin is a plant hormone binding to the plant F-box protein transport inhibitor response 1 (TIR1).^26^ Although no known mammalian targets have been identified for auxin, the indole unit in auxin represents a privileged scaffold that can potentially interact with different proteins.^27, 28^ We reasoned that interrogating the target-binding profiles of these 3 different compounds would yield potential E3 ligases for PROTAC development.

In order to identify the potential targets that these three compounds can bind to in human cells, we designed their corresponding clickable photoaffinity probes **4-6** (Figure 1B). We have employed the clickable photoaffinity probes for target identification of various bioactive compounds.^29-33^ These probes are expected to covalently crosslink with their targets in living cells upon photo-irradiation at 365 nm (Figure 1C). Then the captured targets can be visualized on the gel through in-gel fluorescence scanning and identified through downstream LC-MS/MS analysis. Therefore, the acids were coupled with amine **7** using EDCI as the activating reagent. Then breast cancer MDA-MB-468 cells were treated with different concentrations of the probes, which were then irradiated at 365 nm for 5 min. The lysates were then clicked with a TAMRA-azide (Figure S1) using Cu(I)-catalyzed azide alkyne cycloaddition (aka click chemistry) with TBTA, TCEP and CuSO_4_.^30^ The clicked lysates were then separated on a SDS-PAGE gel for in-gel fluorescence analysis. As shown in Figure 1D, each probe generated dose-dependent labeling of cellular proteins. While these three probes share similar structures, the labeling pattern for each probe is distinct. Strong labeling of prominent bands was observed with probes **4** and **5** while probe **6** provided the least amount of labeling.

To identify proteins that were labeled by these three photoaffinity probes, we clicked the probe-treated cell lysates with a biotin-N_3_ (Figure S1).^30, 31, 33^ Then the biotinylated proteins were enriched using streptavidin-agarose beads. After extensive washings, the bound proteins were eluted for trypsin digestion. The identities of the bound proteins were elucidated using LC-MS/MS analysis (Table S1). The abundance of each protein was quantified by label-free spectra counting. Among the identified proteins, a total of 6 E3 ligases (MARCHF5, RNF121, UBR4, RNF170, HUWE1, SYVN1) were identified for these three probes (Figure 1E). The first four E3 ligases were labeled by all the three probes, suggesting that these E3 ligases might be promiscuous binders of the probes, which share similar structural motifs. While HUWE1 was uniquely labeled by probe **6** and represents a potential E3 ligase target for auxin, in this article, we will focus on synoviolin 1 (SYVN1) that was uniquely identified as a target for probe **4** based on clofibric acid. SYVN1 is known to be a E3 ligase involved in endoplasmic reticulum (ER)-associated degradation (ERAD).^34, 35^ Besides ER localization, SYVN1 has also been known to be localized in the nucleus.^36^

### Validation of SYVN1 as a target of clofibric acid

To validate SYVN1 as a potential target for probe **4**, we first employed biotin-N_3_ click reaction followed by streptavidin pulldown. MDA-MB-468 cells were treated with different concentrations of probe **4** (0, 0.5, 5.0 μM). Then the cells were UV-irradiated at 365 nm for 5 min. The resulting lysates were clicked with a biotin-N_3_ (Figure S1). The biotinylated proteins were then enriched using streptavidin-conjugated agarose beads. After extensive washings, the bound proteins were analyzed by western blot. As shown in Figure 2A, SYVN1 was detected in the streptavidin-bound fraction in a dose-dependent manner. As a control, heat shock protein 90 (Hsp90) was not enriched by the streptavidin pulldown. To further investigate the binding between clofibric acid, we evaluated the effect of clofibramide **8** using a cellular thermal shift assay (CeTSA).^37^ As shown in Figure 2B, clofibramide **8** protected SYVN1 from heat-induced aggregation in MDA-MB-468 cells, supporting that amide **8** binds to SYVN1 in the cells.

**Figure 2.**
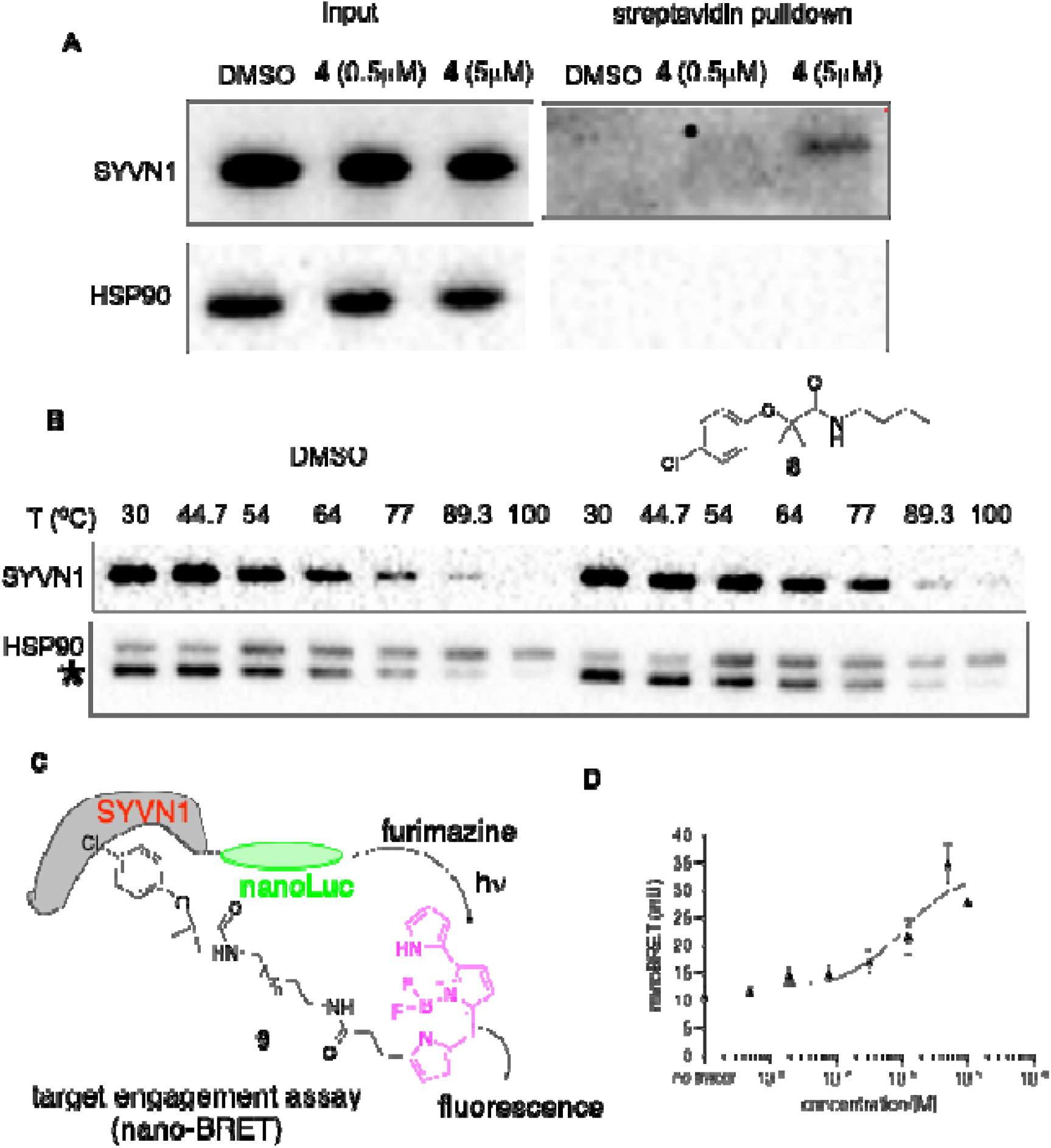
Validation of clofibric acid as a SYVN1 ligand. (A) Probe **4** binds to SYVN1 in MDA-MB-468 cells. The cells were treated with different concentrations of **4** for photocrosslinking. Then the lysates were clicked with a biotin-N_3_ for streptavidin pulldown. The bound proteins were eluted and separated on an SDS-PAGE followed by blotting with indicated antibodies. (B) CeTSA analysis. MDA-MB-468 cells were treated with 0 or 5 μM amide **8**. Then the cells were heated at indicated temperatures for 3 min. The extractable proteins were analyzed by western blotting. Asterisk indicates unstripped SYVN1 in the Hsp90 blot. (C) A schematic drawing of the nano-BRET assay between tracer **9** and SYVN1. (D) Nano-BRET assay between **9** and SYVN1. HEK293T cells were transfected with fusion SYVN1-nanoLuc. Then the cells were treated with increasing concentrations of tracer **9**. To measure the nano-BRET signals, the cells were treated with both nanoLuc substrate furimazine and extracellular nanoLuc inhibitor. The nano-BRET signal was calculated by dividing the light emission from BODIPY between 610-700 nm to the nanoLuc emission between 445-470 nm.

To further investigate the binding between clofibric acid and SYVN1, we designed a cellular nano-BRET (bioluminescence resonance energy transfer) assay.^38, 39^ We created an in-frame fusion between nano-luciferase (nanoLuc) and SYVN1. A small molecule fluorescent tracer **9** (Figure 2C) was also designed and synthesized. Direct binding between clorfibric acid and SYVN1 would be anticipated to bring the BODIPY moiety in probe **9** in close proximity to nanoLuc. As a consequence, a BRET between nanoLuc and BODIPY was expected upon the addition of nanoLuc substrate furimazine (Figure 2C). HEK293T cells were transfected with SYVN1-nanoLuc. Then the cells were treated with a series of concentrations of tracer **9**. The relative nano-BRET signal was measured by dividing the light emission from BODIPY between 610-700 nm to the nanoLuc emission between 445-470 nm. As shown in Figure 2D, titration of tracer **9** into the cells expressing SYVN1-nanoLuc generated a dose-dependent increase of nano-BRET signal with an EC_50_ of 1.07 μM. These data strongly support that clofibric acid and its derivative amide can bind to SYVN1 in live cells, suggesting that clofibric acid can be potentially harnessed for developing novel PROTACs.

### Development of PROTACs based on clofibric acid

With our data demonstrating that clofibric acid can engage SYVN1 in live cells, we were interested in investigating if it could be harnessed for developing PROTACs. SYVN1 has not been employed for developing PROTACs. To test this hypothesis, a series of conjugates between bromodomain containing 4 (BRD4) binder JQ1 and clofibric acid were designed and synthesized. The first series of conjugates **10-16** (Scheme 1) were designed to explore the flexible carbon-based linkers between JQ1 and clofibric acid with different chain length. Additionally, a conformationally more rigid PROTAC **27** was also designed by incorporating a phenylacetylene unit in the linker. Conjugates **10-16** were synthesized from commercially available alkyl diamines (see Supplementary Information), JQ1 acid **26** and clofibric acid (**1**). The diamines were first mono-Boc protected for conjugation with **1** or **26** followed by Boc-deprotection and another amide formation reaction with acid **26** or **1**. Compound **27** was synthesized according to Scheme 1. Amine **17** was coupled to **1** to provide **18**. Sonogashira coupling between alkyne **20** and **18** gave **21**, whose TBS was deprotected using TBAF to generate alcohol **22**, which was then converted to amine **25** through tosylation, azide displacement and Staudinger reaction. Finally, amine **25** was coupled with JQ1 acid **26** to deliver final PROTAC compound **27**.

**Scheme 1.**
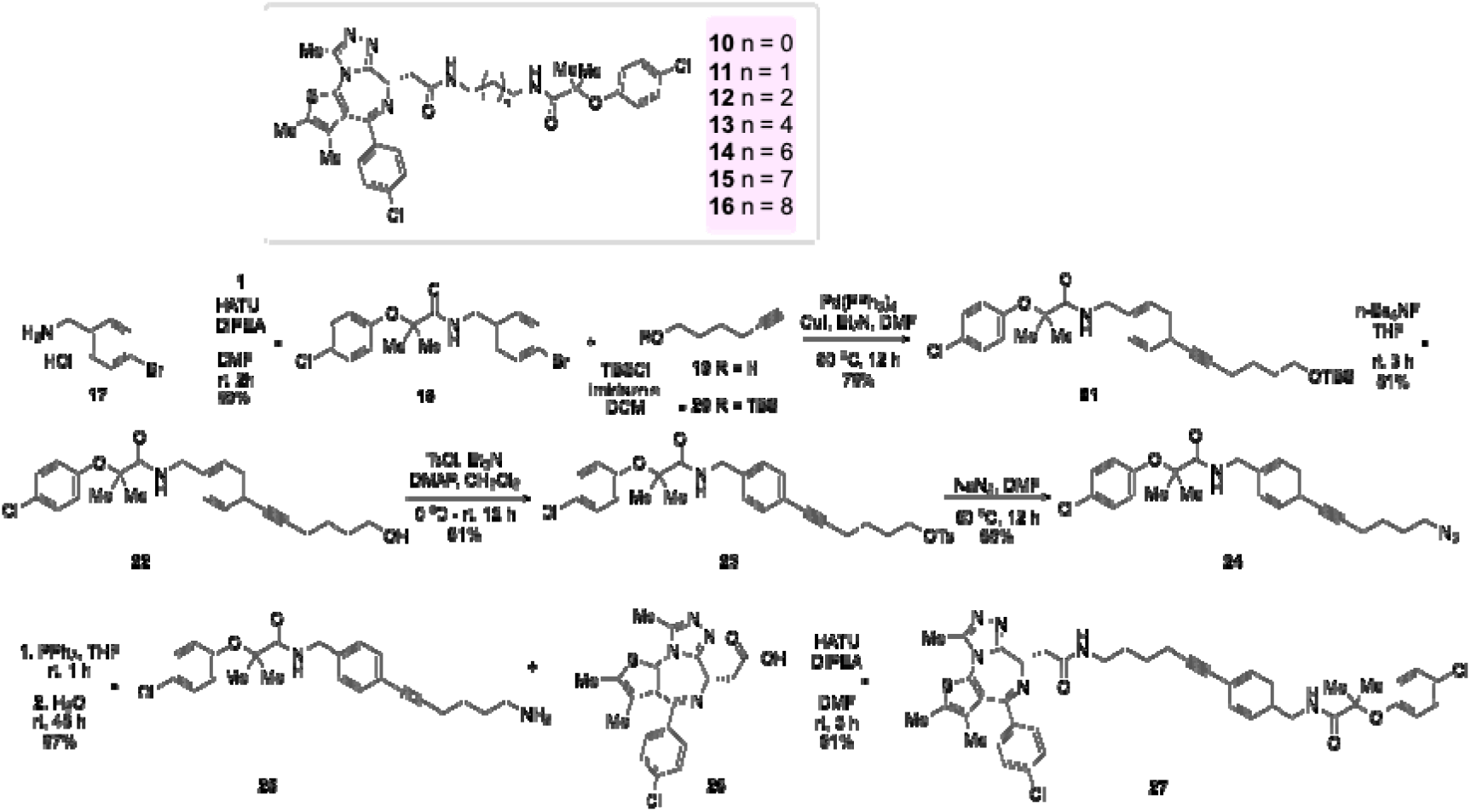
Synthesis of PROTACs. See supplementary information for the detailed synthesis.

To evaluate the potential efficiencies of the newly synthesized PROTACs for BRD4 degradation, we employed CRISPR/Cas9 system to knock-in an 11-amino acid luminescent peptide called HiBit tag into the N-terminus of BRD4 in HEK293T cells.^40^ The level of HiBit-BRD4 in the cells can be conveniently quantified using the nanoLuc complementation assay with LgBit peptide and nanoLuc substrate (Promega). To confirm that the engineered HEK293T cells were responsive to BRD4 degradation by a PROTAC, we treated the cells with dBET6,^41^ a PROTAC based on JQ1 and a CRBN ligand. As shown in Figure S2, dBET6 efficiently degraded BRD4 with an IC_50_ of 43.6 nM. Therefore, we screened the newly synthesized PROTACs for potential BRD4 degradation using the HiBit-BRD4 system. Compounds **10-16** were first evaluated at 5 and 20 μM in the HEK293T cells with HiBit-BRD4 for 20 h. As shown in Figure 3A, the compounds presented varied BRD4 degradation efficiency. Higher degradation was observed at 20 μM versus 5 μM for all the compounds, supporting their dose-dependency. At 20 μM, higher degradation efficiency was observed with increased chain length from 2 to 9 carbons (cmpd **10-15**). Further increase of the chain length to 10 carbons (i.e. cpmd **16**) did not yield further increase of BRD4 degradation. Besides the linker length, the linker composition also plays an important role in the target degradation efficiency of PROTACs.^2^ We therefore designed a conformationally-constrained PROTAC **27** (Scheme 1) by incorporating a phenylacetylene unit into the linker while keeping the linker length to be 9-10 carbon atoms. Gratifyingly, cmpd **27** showed significantly enhanced BRD4 degradation compared to **10-16** (Figure 3A). We further evaluated BRD4 degradation by **27** using different doses. As shown in Figure 3B, **27** produced dose-dependent degradation of BRD4 with an IC_50_ of 0.93 μM in the HiBit-BRD4 assay. The BRD4 degradation by **27** was further confirmed in native HEK293T cells using western blot assay. In this case, the endogenous BRD4 was also significantly degraded by **27**. Significant degradation was observed even at 100 nM (Figure 3C). Together, these results support that clofibric acid can be utilized as a novel small molecule ligand (MW = 214) for the design of PROTACs.

**Figure 3.**
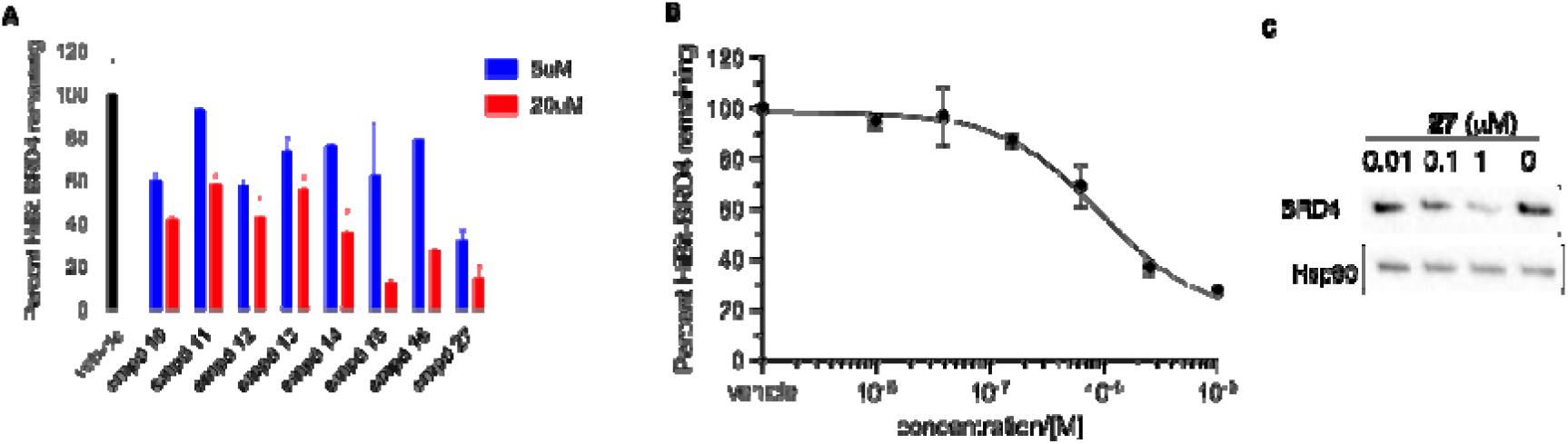
BRD4 degradation by PROTACs **10-27**. (A) HiBit-BRD4 degradation. HEK293T cells with HiBit-BRD4 were treated with 5 or 20 μM of different compounds for 20 h. Then the remaining level of HiBit-BRD4 was quantified using a lytic nano-Glo^®^ assay kit (Promega). (B) Dose-dependent degradation of HiBit-BRD4 by **27**. HEK293T cells with HiBit-BRD4 were treated with different concentrations of **27** for 20 h. Then the remaining level of HiBit-BRD4 was quantified using a lytic nano-Glo^®^ assay kit (Promega). The IC_50_ was calculated by non-linear regression analysis in Prism 10. (C) Cmpd **27** induced degradation of endogenous BRD4. HEK293T cells were treated with different concentrations of **27** for 20 h. Then the lysates were prepared for western blot analysis with indicated antibodies. Hsp90 was used as a loading control.

### PROTAC 27 degraded BRD4 by a PROTAC mechanism

Having identified **27** to be able to degrade BRD4, we investigated its mechanism of action to ascertain that the induced BRD4 degradation was through a PROTAC mechanism. First, we asked if a ternary complex involving BRD4 and SYVN1 could be induced in the presence of **27**. To this end, we expressed Flag-tagged BRD4 in HEK 293T cells. Then the lysates were subjected to co-immunoprecipitation (co-IP) using anti-Flag (M2) in the presence or absence of **27**. In the absence of **27**, BRD4 and SYVN1 did not form a complex (Figure 4A). Upon the addition of **27**, we observed SYVN1 in the BRD4 immune complex, suggesting that **27** induced ternary complex formation between BRD4 and SYVN1 to promote BRD4 degradation. Next, we investigated if BRD4 degradation induced by **27** was mediated by the UPS. BRD4 degradation was observed 2 h post treatment with **27** in HEK 293T cells. Co-treating the cells with a proteasome inhibitor MG132 rescued the degradation induced by cmpd **27**, demonstrating the requirement of intact proteasome activity to induce BRD4 degradation.

**Figure 4.**
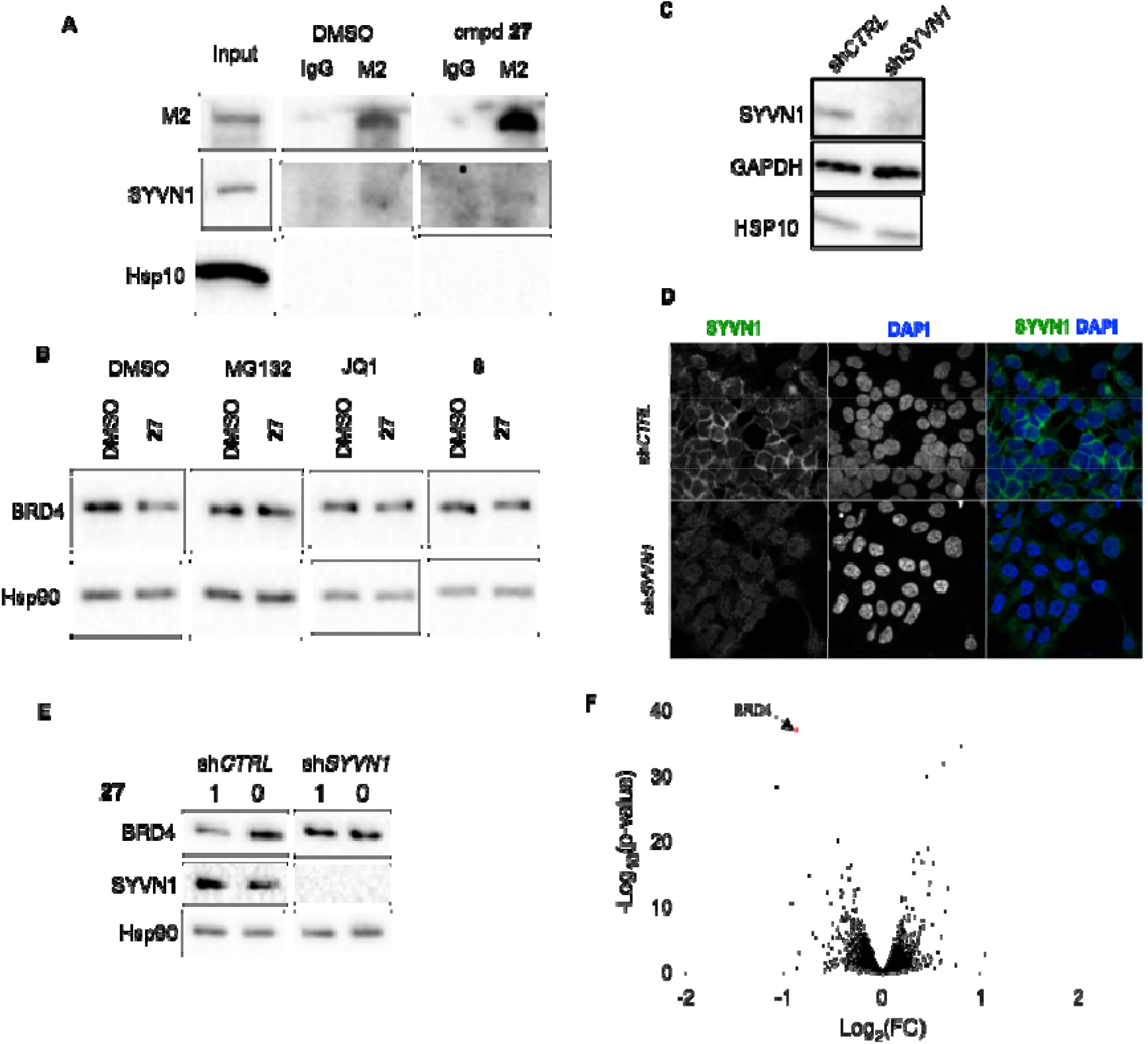
PROTAC **27** selectively degraded BRD4. (A) Cmpd **27** induced ternary complex formation between BRD4 and SYVN1. HEK 293T cells were transfected with Flag-tagged BRD4. Then the lysates were subjected to co-IP with anti-Flag in the presence of 0 or 1 μM cmpd **27**. The bound immune complexes were then analyzed by western blot with indicated antibodies. (B) Cmpd **27** degraded BRD4 through a PROTAC mechanism. HEK 293T cells were treated with DMSO or **27** (1 μM) along with indicated drugs (MG132 (5 μM), JQ1 (5 μM), cmpd **8** (5 μM)) for 2 h. Then the lysates were analyzed by western blot with indicated antibodies. (C) sh*RNA* knockdown of SYVN1. HEK 293T cells were transduced with lentiviruses expressing sh*CTRL* or sh*SYVN1*. Then the lysates from each cell lines were analyzed by western blot using indicated antibodies. (D) Validation of SYVN1 knockdown in HEK 293T cells by immunofluorescence. HEK 293T cells were sh*CTRL* or sh*SYVN1* were analyzed by immunofluorescence. The cells were fixed, permeabilized and stained with anti-SYVN1. The nuclei were counter stained with DAPI. (E) Cmpd **27**-induced degradation of BRD4 required SYVN1. HEK 293T cells were transduced with lentiviruses expressing sh*CTRL* or sh*SYVN1*. Then the cells were treated with 0 or 1 μM cmpd **27** for 2 h. The lysates were analyzed by western blot with indicated antibodies. (F) Quantitative tandem mass tagging (TMT)-based proteomic profiling of cmpd **27** in HEK 293T cells. HEK 293 cells were treated with 0 or 1 μM cmpd **27** for 20 h. Then the lysates were prepared for TMT-based LC-MS/MS analysis. The data were presented as a volcano plot. BRD4 is highlighted in red. The full proteomics data can be found in Table S2.

PROTACs degrade target proteins through the formation of ternary complexes. Thus, we conducted competition experiments to assess the BRD4 degradation by **27**. Addition of JQ1 to the cells treated with **27** abolished BRD4 degradation (Figure 4B). Similarly, addition of clofibramide **8** also inhibited **27**-induced BRD4 degradation (Figure 4B). These data support the critical importance of functional ternary complex formation to induce BRD4 degradation by **27**. To further investigate the role of SYVN1 in mediating **27**-induced BRD4 degradation, we employed lentiviral sh*RNA* to knockdown the expression of SYVN1. As shown in Figure 4C, SYVN1 protein expression was significantly knocked down using sh*SYVN1*. The knockdown was further validated by immunofluorescence using confocal microscopy (Figure 4D). Consistent with previous findings that SYVN1 was found to be localized in nucleus besides cytosol,^36^ we also found that SYVN1 was present in the nucleus in HEK 293T cells (Figure 4D). When the cells with sh*SYVN1* were treated with **27**, BRD4 degradation was not observed (Figure 4E). On the other hand, the sh*CTRL* cells showed efficient BRD4 degradation, similar to the un-transduced cells shown in Figure 4B. These results demonstrated that BRD4 degradation by **27** was SYVN1-dependent. Finally, the global protein degradation selectivity of **27** was evaluated using tandem mass tag (TMT)-based quantitative proteomics. HEK 293T cells were treated with **27** for 20 h. Then the lysates were prepared and digested with trypsin. Equal amounts of peptides from each sample were labeled with TMT reagents. Then the samples were analyzed by LC-MS/MS. As shown in Figure 4F, BRD4 was selectively and significantly degraded among the 6,600 proteins quantified. Altogether, these results demonstrate cmpd **27** is a *bona fide* PROTAC to selectively degrade BRD4 through the engagement of SYVN1.

## CONCLUSIONS

PROTACs represent a new therapeutic modality for numerous targets, many of which are undrugged yet.^42^ While PROTACs are generally higher in molecular weight, many of the clinical candidates show reasonable oral bioavailability and are being evaluated as oral medications.^22^ The human genome encodes ∼600 E3 ligases, yet only a few have been successfully harnessed for developing PROTACs. The most commonly used ones are CRBN and VHL. Therefore, expanding the druggable E3 ligases for developing PROTACs has a tremendous potential to further expand the druggable genome. Furthermore, increasing the targetable E3 ligases for developing PROTACs can offer potential target selectivity and safety. With a goal of identifying small fragment-like ligands for E3 ligases, we resorted to clinically approved drugs and endogenous bioactive compounds as they would be expected to have fewer undesirable effects. Using clickable photoaffinity probes derived from clofibric acid, naproxen and auxin, we identified a number of E3 ligases that can be potential novel off targets of these carboxylic acids. Among these, SYVN1 was specifically identified as a novel E3 ligase target for clofibric acid. Clofibric acid demonstrated engagement with SYVN1 in human cells using both CeTSA and nano-BRET assay. As a small molecule with a MW of only 214 Daltons, we designed a series of potential PROTACs based on BRD4 ligand JQ1. Gratifyingly, all the synthesized PROTACs exhibited some degree of BRD4 degradation. The PROTACs showed strong dependence on the linker length and composition. The more rigid linker in cmpd **27** demonstrated the highest potency in BRD4 degradation. The BRD4 degradation induced by **27** was dependent on the functionally intact UPS and ternary complex formation. Finally, PROTAC **27** demonstrated high selectivity toward BRD4 in the human proteome as assessed by TMT-based quantitative proteomics. HUWE1 was identified as a potential target for auxin. Validation of these results and its application in the PROTAC design are ongoing and will be reported in due course. In conclusion, our results support the utility of clofibric acid as a novel SYVN1 ligand for development of PROTACs. The small molecular weight of clofibric acid is an appealing feature for future design of PROTACs as candidates for both clinical development and tools for biomedical research.

## Supporting Information Available

Additional data and experimental details are presented in the supplementary information.

## Acknowledgement

We greatly appreciate the financial supports provided from National Institutes of Health (R01GM122820) and Oregon Health & Science University (OHSU). We thank OHSU Biophysical Shared Resources Core for providing various biophysical instrument to support this work. We appreciate OHSU Integrated genomics core for authenticating the cell lines through STR profiling. Mass spectrometric analysis was performed by the OHSU Proteomics Shared Resource (RRID: SCR_009991) with partial support from NIH core grants P30EY010572, P30CA069533, S10 OD028533 and OHSU Emerging Technology Fund.

## References

1. Schapira, M. Calabrese, M. F. Bullock, A. N. Crews, C. M. Targeted protein degradation: expanding the toolbox. Nature reviews. Drug discovery 2019, 18 (12), 949–963, 10.1038/s41573-019-0047-y.

2. Martín-Acosta, P. Xiao, X. PROTACs to address the challenges facing small molecule inhibitors. Eur. J. Med. Chem. 2021, 210, 112993, 10.1016/j.ejmech.2020.112993.

3. Schwalm, M. P. Krämer, A. Dölle, A. Weckesser, J. Yu, X. Jin, J. Saxena, K. Knapp, S. Tracking the PROTAC degradation pathway in living cells highlights the importance of ternary complex measurement for PROTAC optimization. Cell Chem. Biol. 2023, 30 (7), 753–765.e8, 10.1016/j.chembiol.2023.06.002.

4. Li, K. Crews, C. M. PROTACs: past, present and future. Chem Soc Rev 2022, 51 (12), 5214–5236, 10.1039/d2cs00193d.

5. Békés, M. Langley, D. R. Crews, C. M. PROTAC targeted protein degraders: the past is prologue. Nature reviews. Drug discovery 2022, 21 (3), 181–200, 10.1038/s41573-021-00371-6.

6. Wang, X. Li, Y. He, M. Kong, X. Jiang, P. Liu, X. Diao, L. Zhang, X. Li, H. Ling, X. Xia, S. Liu, Z. Liu, Y. Cui, C. P. Wang, Y. Tang, L. Zhang, L. He, F. Li, D. UbiBrowser 2.0: a comprehensive resource for proteome-wide known and predicted ubiquitin ligase/deubiquitinase-substrate interactions in eukaryotic species. Nucleic Acids Res. 2022, 50 (D1), D719–d728, 10.1093/nar/gkab962.

7. Ge, J. Li, S. Weng, G. Wang, H. Fang, M. Sun, H. Deng, Y. Hsieh, C. Y. Li, D. Hou, T. PROTAC-DB 3.0: an updated database of PROTACs with extended pharmacokinetic parameters. Nucleic Acids Res. 2025, 53 (D1), D1510–d1515, 10.1093/nar/gkae768.

8. Ishida, T. Ciulli, A. E3 Ligase Ligands for PROTACs: How They Were Found and How to Discover New Ones. SLAS Discov 2021, 26 (4), 484–502, 10.1177/2472555220965528.

9. Gui, W. Goss, A. Kodadek, T. A Functional Assay for Mining Noninhibitory Enzyme Ligands from One Bead One Compound Libraries: Application to E3 Ubiquitin Ligases. J. Am. Chem. Soc. 2025, 147 (35), 31630–31638, 10.1021/jacs.5c07307.

10. Lu, G. Middleton, R. E. Sun, H. Naniong, M. Ott, C. J. Mitsiades, C. S. Wong, K. K. Bradner, J. E. Kaelin, W. G., Jr. The myeloma drug lenalidomide promotes the cereblon-dependent destruction of Ikaros proteins. Science 2014, 343 (6168), 305–9, 10.1126/science.1244917.

11. Buckley, D. L. Van Molle, I. Gareiss, P. C. Tae, H. S. Michel, J. Noblin, D. J. Jorgensen, W. L. Ciulli, A. Crews, C. M. Targeting the von Hippel-Lindau E3 ubiquitin ligase using small molecules to disrupt the VHL/HIF-1α interaction. J. Am. Chem. Soc. 2012, 134 (10), 4465–8, 10.1021/ja209924v.

12. Ohoka, N. Okuhira, K. Ito, M. Nagai, K. Shibata, N. Hattori, T. Ujikawa, O. Shimokawa, K. Sano, O. Koyama, R. Fujita, H. Teratani, M. Matsumoto, H. Imaeda, Y. Nara, H. Cho, N. Naito, M. In Vivo Knockdown of Pathogenic Proteins via Specific and Nongenetic Inhibitor of Apoptosis Protein (IAP)-dependent Protein Erasers (SNIPERs). J. Biol. Chem. 2017, 292 (11), 4556–4570, 10.1074/jbc.M116.768853.

13. Schröder, M. Renatus, M. Liang, X. Meili, F. Zoller, T. Ferrand, S. Gauter, F. Li, X. Sigoillot, F. Gleim, S. Stachyra, T.-M. Thomas, J. R. Begue, D. Khoshouei, M. Lefeuvre, P. Andraos-Rey, R. Chung, B. Ma, R. Pinch, B. Hofmann, A. Schirle, M. Schmiedeberg, N. Imbach, P. Gorses, D. Calkins, K. Bauer-Probst, B. Maschlej, M. Niederst, M. Maher, R. Henault, M. Alford, J. Ahrne, E. Tordella, L. Hollingworth, G. Thomä, N. H. Vulpetti, A. Radimerski, T. Holzer, P. Carbonneau, S. Thoma, C. R. DCAF1-based PROTACs with activity against clinically validated targets overcoming intrinsic- and acquired-degrader resistance. Nature communications 2024, 15 (1), 275, 10.1038/s41467-023-44237-4.

14. Zhang, X. Luukkonen, L. M. Eissler, C. L. Crowley, V. M. Yamashita, Y. Schafroth, M. A. Kikuchi, S. Weinstein, D. S. Symons, K. T. Nordin, B. E. Rodriguez, J. L. Wucherpfennig, T. G. Bauer, L. G. Dix, M. M. Stamos, D. Kinsella, T. M. Simon, G. M. Baltgalvis, K. A. Cravatt, B. F. DCAF11 Supports Targeted Protein Degradation by Electrophilic Proteolysis-Targeting Chimeras. J. Am. Chem. Soc. 2021, 143 (13), 5141–5149, 10.1021/jacs.1c00990.

15. Zhang, X. Crowley, V. M. Wucherpfennig, T. G. Dix, M. M. Cravatt, B. F. Electrophilic PROTACs that degrade nuclear proteins by engaging DCAF16. Nat. Chem. Biol. 2019, 15 (7), 737–746, 10.1038/s41589-019-0279-5.

16. Spradlin, J. N. Hu, X. Ward, C. C. Brittain, S. M. Jones, M. D. Ou, L. To, M. Proudfoot, A. Ornelas, E. Woldegiorgis, M. Olzmann, J. A. Bussiere, D. E. Thomas, J. R. Tallarico, J. A. McKenna, J. M. Schirle, M. Maimone, T. J. Nomura, D. K. Harnessing the anti-cancer natural product nimbolide for targeted protein degradation. Nat. Chem. Biol. 2019, 15 (7), 747–755, 10.1038/s41589-019-0304-8.

17. Schneekloth, A. R. Pucheault, M. Tae, H. S. Crews, C. M. Targeted intracellular protein degradation induced by a small molecule: En route to chemical proteomics. Bioorg. Med. Chem. Lett. 2008, 18 (22), 5904–8, 10.1016/j.bmcl.2008.07.114.

18. Henning, N. J. Manford, A. G. Spradlin, J. N. Brittain, S. M. Zhang, E. McKenna, J. M. Tallarico, J. A. Schirle, M. Rape, M. Nomura, D. K. Discovery of a Covalent FEM1B Recruiter for Targeted Protein Degradation Applications. J. Am. Chem. Soc. 2022, 144 (2), 701–708, 10.1021/jacs.1c03980.

19. Basu, A. A. Zhang, C. Riha, I. A. Magassa, A. Campos, M. A. Caldwell, A. G. Ko, F. Zhang, X. A CRISPR activation screen identifies FBXO22 supporting targeted protein degradation. Nat. Chem. Biol. 2024, 20 (12), 1608–1616, 10.1038/s41589-024-01655-9.

20. Poirson, J. Cho, H. Dhillon, A. Haider, S. Imrit, A. Z. Lam, M. H. Y. Alerasool, N. Lacoste, J. Mizan, L. Wong, C. Gingras, A. C. Schramek, D. Taipale, M. Proteome-scale discovery of protein degradation and stabilization effectors. Nature 2024, 628 (8009), 878–886, 10.1038/s41586-024-07224-3.

21. Khan, S. Zhang, X. Lv, D. Zhang, Q. He, Y. Zhang, P. Liu, X. Thummuri, D. Yuan, Y. Wiegand, J. S. Pei, J. Zhang, W. Sharma, A. McCurdy, C. R. Kuruvilla, V. M. Baran, N. Ferrando, A. A. Kim, Y. M. Rogojina, A. Houghton, P. J. Huang, G. Hromas, R. Konopleva, M. Zheng, G. Zhou, D. A selective BCL-X(L) PROTAC degrader achieves safe and potent antitumor activity. Nat. Med. 2019, 25 (12), 1938–1947, 10.1038/s41591-019-0668-z.

22. Srivastava, A. Pike, A. Swedrowska, M. Nash, S. Grime, K. In Vitro ADME Profiling of PROTACs: Successes, Challenges, and Lessons Learned from Analysis of Clinical PROTACs from a Diverse Physicochemical Space. J. Med. Chem. 2025, 68 (9), 9584–9593, 10.1021/acs.jmedchem.5c00358.

23. Issemann, I. Green, S. Activation of a member of the steroid hormone receptor superfamily by peroxisome proliferators. Nature 1990, 347 (6294), 645–50, 10.1038/347645a0.

24. Forman, B. M. Chen, J. Evans, R. M. Hypolipidemic drugs, polyunsaturated fatty acids, and eicosanoids are ligands for peroxisome proliferator-activated receptors alpha and delta. Proc. Natl. Acad. Sci. U. S. A. 1997, 94 (9), 4312–7, 10.1073/pnas.94.9.4312.

25. Duggan, K. C. Hermanson, D. J. Musee, J. Prusakiewicz, J. J. Scheib, J. L. Carter, B. D. Banerjee, S. Oates, J. A. Marnett, L. J. (R)-Profens are substrate-selective inhibitors of endocannabinoid oxygenation by COX-2. Nat. Chem. Biol. 2011, 7 (11), 803–9, 10.1038/nchembio.663.

26. Tan, X. Calderon-Villalobos, L. I. Sharon, M. Zheng, C. Robinson, C. V. Estelle, M. Zheng, N. Mechanism of auxin perception by the TIR1 ubiquitin ligase. Nature 2007, 446 (7136), 640–5, 10.1038/nature05731.

27. Chao, B. Li, B. X. Xiao, X. The chemistry and pharmacology of privileged pyrroloquinazolines. MedChemComm 2015, 6 (4), 510–520.

28. Welsch, M. E. Snyder, S. A. Stockwell, B. R. Privileged scaffolds for library design and drug discovery. Curr. Opin. Chem. Biol. 2010, 14 (3), 347–361, 10.1016/j.cbpa.2010.02.018.

29. Li, B. X. Chen, J. Chao, B. David, L. L. Xiao, X. Anticancer pyrroloquinazoline LBL1 targets nuclear lamins. ACS Chem. Biol. 2018, 13, 1380–1387, 10.1021/acschembio.8b00266.

30. Xiao, X. Li, B. X. Identification of lamins as the molecular targets of LBL1 using a clickable photoaffinity probe. Methods Enzymol. 2020, 633, 185–201, 10.1016/bs.mie.2019.02.038.

31. Warren, J. Wang, J. Dhoro, F. Chao, B. Reddy, A. Petrie, S. K. David, L. L. Xiao, X. Li, B. X. SMAP3-ID for Identification of Endogenous Protein-Protein Interactions Reveals Regulation of Mitochondrial Activity by Lamins. JACS Au 2025, 5 (1), 302–319, 10.1021/jacsau.4c00988.

32. Warren, J. Wang, J. Li, B. X. Xiao, X. Design, synthesis and evaluation of clickable photoaffinity probes for nuclear lamins. Bioorg. Med. Chem. Lett. 2025, 129, 130392, 10.1016/j.bmcl.2025.130392.

33. Martín-Acosta, P. Meng, Q. Klimek, J. Reddy, A. P. David, L. Petrie, S. K. Li, B. X. Xiao, X. A clickable photoaffinity probe of betulinic acid identifies tropomyosin as a target. Acta Pharm Sin B 2022, 12 (5), 2406–2416, 10.1016/j.apsb.2021.12.008.

34. Amano, T. Yamasaki, S. Yagishita, N. Tsuchimochi, K. Shin, H. Kawahara, K. Aratani, S. Fujita, H. Zhang, L. Ikeda, R. Fujii, R. Miura, N. Komiya, S. Nishioka, K. Maruyama, I. Fukamizu, A. Nakajima, T. Synoviolin/Hrd1, an E3 ubiquitin ligase, as a novel pathogenic factor for arthropathy. Genes Dev. 2003, 17 (19), 2436–49, 10.1101/gad.1096603.

35. Kikkert, M. Doolman, R. Dai, M. Avner, R. Hassink, G. van Voorden, S. Thanedar, S. Roitelman, J. Chau, V. Wiertz, E. Human HRD1 is an E3 ubiquitin ligase involved in degradation of proteins from the endoplasmic reticulum. J. Biol. Chem. 2004, 279 (5), 3525–34, 10.1074/jbc.M307453200.

36. Leitman, J. Shenkman, M. Gofman, Y. Shtern, N. O. Ben-Tal, N. Hendershot, L. M. Lederkremer, G. Z. Herp coordinates compartmentalization and recruitment of HRD1 and misfolded proteins for ERAD. Mol. Biol. Cell 2014, 25 (7), 1050–60, 10.1091/mbc.E13-06-0350.

37. Martinez Molina, D. Jafari, R. Ignatushchenko, M. Seki, T. Larsson, E. A. Dan, C. Sreekumar, L. Cao, Y. Nordlund, P. Monitoring drug target engagement in cells and tissues using the cellular thermal shift assay. Science 2013, 341 (6141), 84–7, 10.1126/science.1233606.

38. Robers, M. B. Dart, M. L. Woodroofe, C. C. Zimprich, C. A. Kirkland, T. A. Machleidt, T. Kupcho, K. R. Levin, S. Hartnett, J. R. Zimmerman, K. Niles, A. L. Ohana, R. F. Daniels, D. L. Slater, M. Wood, M. G. Cong, M. Cheng, Y. Q. Wood, K. V. Target engagement and drug residence time can be observed in living cells with BRET. Nature communications 2015, 6, 10091, 10.1038/ncomms10091.

39. Monroy, E. Y. Yu, X. Lu, D. Qi, X. Wang, J. One Tracer, Dual Platforms: Unlocking Versatility of Fluorescent Probes in TR-FRET and NanoBRET Target Engagement Assays. ACS Med. Chem. Lett. 2025, 16 (8), 1554–1561, 10.1021/acsmedchemlett.5c00171.

40. Schwinn, M. K. Machleidt, T. Zimmerman, K. Eggers, C. T. Dixon, A. S. Hurst, R. Hall, M. P. Encell, L. P. Binkowski, B. F. Wood, K. V. CRISPR-Mediated Tagging of Endogenous Proteins with a Luminescent Peptide. ACS Chem. Biol. 2018, 13 (2), 467–474, 10.1021/acschembio.7b00549.

41. Winter, G. E. Mayer, A. Buckley, D. L. Erb, M. A. Roderick, J. E. Vittori, S. Reyes, J. M. di Iulio, J. Souza, A. Ott, C. J. Roberts, J. M. Zeid, R. Scott, T. G. Paulk, J. Lachance, K. Olson, C. M. Dastjerdi, S. Bauer, S. Lin, C. Y. Gray, N. S. Kelliher, M. A. Churchman, L. S. Bradner, J. E. BET Bromodomain Proteins Function as Master Transcription Elongation Factors Independent of CDK9 Recruitment. Mol. Cell 2017, 67 (1), 5–18.e19, 10.1016/j.molcel.2017.06.004.

42. Jiang, J. Yuan, J. Hu, Z. Zhang, Y. Zhang, T. Xu, M. Long, M. Fan, Y. Tanyi, J. L. Montone, K. T. Tavana, O. Vonderheide, R. H. Chan, H. M. Hu, X. Zhang, L. Systematic illumination of druggable genes in cancer genomes. Cell reports 2022, 38 (8), 110400, 10.1016/j.celrep.2022.110400.

