## Supplementary information for "Identification of clofibric acid as a SYVN1 ligand for PROTAC development"

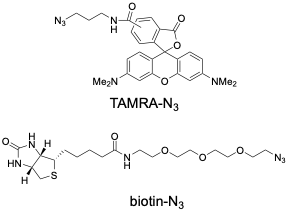

**Figure S1**. Chemical structures of TAMRA-N_3_ and biotin-N_3_ used in this paper.

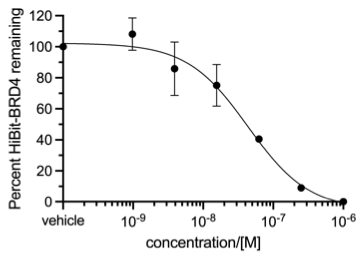

**Figure S2**. dBET6 dose-dependently induced degradation of BRD4 assessed by HiBit-BRD4 assay. HEK29T cells with HiBit-BRD4 were treated with different concentrations of dBET6 for 20 h. Then the remaining HiBit-BRD4 was assessed using Nano-Glo^®^ HiBiT Lytic Detection System (Promega).

**Experimental Section**.

**Cell Lines and Culture.** MDA-MB-468 cells were purchased from Developmental Therapeutics Program at the National Cancer Institute. HEK 293T cells were purchased from American Tissue Culture Collection (ATCC). Cells were cultured in high-glucose Dulbecco’s Modified Eagle’s Medium (DMEM, Life Technologies) supplemented with 10% FBS (Hyclone) and nonessential amino acids (Life Technologies) at 37 °C with 5% CO_2_. All the cells were authenticated using STR profiling and used within 50 passages.

**Plasmids**. Flag-HaloTag-BRD4 was engineered by inserting Flag-tagged HaloTag to the N-terminus of BRD4. The in-frame construct was generated by gene synthesis (Genewiz, South Plainfield, NJ) and subcloned into the third-generation lentiviral vector pLenti6.2 (Addgene). A nanoLuc–SYVN1 fusion was engineered by inserting nanoLuc to the C-terminus of SYVN1. The in-frame construct was generated by gene synthesis (Genewiz, South Plainfield, NJ) and subsequently subcloned into the third-generation lentiviral vector pLenti6.2/V5-DEST-SYVN1 (DNASU Plasmid Repository). The sequences of the final plasmids were confirmed by next-generation sequencing.

**Antibodies**. The following primary antibodies were used for western blotting: anti-LA/C (mouse, Sigma, Catalog No. SAB4200236, 1:2000), anti-BRD4 (mouse, Cell signaling Technology, Catalog No. 63759, 1:1000), anti-HSP10 (rabbit, Bethyl Laboratories, Catalog. No A304-842A-M, 1:1000), anti-Hsp90 (rabbit, Cell Signaling Technology, Catalog No. 4874, 1:1000, anti-GAPDH (mouse, Santa Cruz Biotechnology, Catalog No. sc-32233, 1:4000), anti-Flag (M2, mouse, Sigma, F3165, 1:4000), anti-SYVN1 (rabbit, Bethyl Laboratories, Catalog No. A302-946A, (1:2000). The HRP-conjugated secondary antibodies (anti-mouse and anti-rabbit) were from Jackson ImmunoResearch Laboratories and used at 1:4000 in 5% non-fat dry milk in TBST.

**Western blot**. The lysates or eluates from streptavidin pulldown were separated on 4-20% TGX precast SDS-PAGE gels (Bio-Rad). Upon electrophoresis, the proteins were electro-transferred to nitrocellulose membranes (Bio-Rad). The membranes were then blocked in 5% non-fat dry milk for 1 h at room temperature. Then the membranes were incubated with primary antibodies at 4 ^o^C overnight. The membranes were then washed 3x with TBST and incubated with HRP-conjugated secondary antibodies (Jackson ImmunoResearch Laboratories, 1:4000) in 5% nonfat dry milk in TBST for 1 h at room temperature. Images were captured using Clarity Western ECL Substrate (Bio-Rad) in ChemiDoc^MP^ (Bio-Rad).

**Nano-BRET assay**. HEK 293T cells were plated into a 24-well plate. The cells were attached to the bottom for overnight. Then the cells were transfected with 50 ng nanoLuc-SYVN1 using Lipofectamine^TM^ 2000 (Thermo Fisher) according to the manufacturer’s instructions. Six hours later, the transfected cells were replated into a white opaque-bottom 96-well plate. The cells were allowed to attach the bottom for overnight. The cells were then treated with tracer compound 9 for 2 h. To measure the nano-BRET signal in the cells, the Intracellular TE Nano-Glo^®^ substrate/inhibitor system (Promega) was used by following the manufacturer’s instructions. Briefly, the plate was equilibrated to room temperature. Then a 3x stock of nanoLuc substrate and extracellular nanoLuc inhibitor in Opti-MEM (Thermo Fisher) was added to each well. The plate was shaken on an orbital shaker for 15 min at room temperature. The luminescence values at 610-700 nm and 445-470 nm were then measured on a Tecan Plate Reader Spark with an integration time of 500 ms. The nanoBRET values were calculated by dividing the readings at 610-700 nm to those at 445-470 nm, which was then multiplied by 1,000.

**Co-IP assay**. A 10-cm plate of HEK 293T cells were transfected with 4 μg of Flag-HaloTag-BRD4 using Lipofectamine^TM^ 2000 (Thermo Fisher) according to the manufacturer’s instructions. Twenty-four hours later, the cells were harvested by scaping and centrifuged at 200x g for 5 min at 4 ^o^C. The cell pellets were washed twice with ice-cold PBS. Then the cells were lysed in 1 mL of lysis buffer (50 mM TrisHCl, 5 mM EDTA, 150 mM NaCl, 1 mM DTT, 0.5% Nonidet P-40, pH 8.0) supplemented with 1 mM PMSF and protease inhibitor cocktail. The lysates were cleared by centrifuging at 14,000x rpm for 15 min on a tabletop centrifuge at 4 ^o^C. The supernatant was precleared with 2 μg mouse IgG (Jackson ImmunoResearch) and 20 μL protein A/G beads (Pierce) by tumbling at 4 ^o^C for 2 h. The precleared lysates were then equally divided two halves. One half was treated with DMSO while the other half was treated with 1 μM cmpd **27**. The treated lysates were then mixed with either 1 μg mouse IgG or anti-Flag (M2, Sigma) for overnight 4 ^o^C. Then 10 μL of protein A/G beads was added to each sample. The samples were tumbled at 4 ^o^C for 1 h. The unbound proteins were removed by centrifuging at 3,000x rpm for 2 min on a tabletop centrifuge at 4 oC and the beads were washed with 3x500 μL of lysis buffer. The bound proteins were eluted with 1x SDS-PAGE buffer (Bio-rad) supplemented with 5% beta-mercaptoethanol by heating at 95 ^o^C for 5 min. The eluted proteins were analyzed by western blotting.

**Generation of HiBit-BRD4 cells**. Introduction of a HiBiT coding sequence into the endogenous *BRD4* locus in HEK 293T cells was done via CRISPR-Cas9 genome editing (Integrated DNA Technologies, IDT). The HDR duplex template and gRNA were resuspended with 20 µL nuclease-free duplex buffer (IDT) to get 100 μM stock and stored at -20 ºC. To prepare 1 µM working stocks, 0.5 µl of 100 μM stock solution was diluted with 49.5 µl nuclease-free duplex buffer and stored at -20 ºC. Three µL of each 1 μM HDR solution was heated for 5 minutes at 95 ºC. After heating, the complex was gradually cooled to room temperature. The oligo complex was then incubated at room temperature for 5 minutes with the HiFi Cas9 Nuclease (IDT) to form the ribonucleoprotein (RNP) complex. The CRISPR-Cas9 RNP complex was transfected with Lipofectamine CRISPRMAZ Cas9 Transfection Reagent (ThermoFisher Scientfic) according to manufacturer’s instructions. The edited polyclonal cells were used directly for HiBit assay. The oligo sequences are below:

>gRNA sequence

ACTAGCATGTCTGCGGAGAG

>HDR donor sequence (+)

CATTACTGGCAGATTTCTCAATCTCGTCCCAGGGCCGCTCTCCGCAGAGCTAATCTTCTTGAACAGCCGCCAGCCGCTCACCATGCTAGTGATCCCATCACATTCTTCACCAGGCACTCTA

>HDR donor sequence (-)

TAGAGTGCCTGGTGAAGAATGTGATGGGATCACTAGCATGGTGAGCGGCTGGCGGCTGTTCAAGAAGATTAGCTCTGCGGAGAGCGGCCCTGGGACGAGATTGAGAAATCTGCCAGTAATG

**HiBit-BRD4 degradation**. HEK 293T cells with HiBit-BRD4 were plated into white opaque bottom 96-well plate at 10,000 cells/100 μL/well. The cells were treated with different concentrations of compounds for 20 h at 37 ^o^C. The plate was brought to room temperature and 50 μL of the culture media was removed. Then 60 μL of the reconstituted lytic nano-Glo^®^ HiBit assay mixture (Promega) was added to each well. The plate was gently shaken on a orbital shaker for 15 min at room temperature. Then the remaining luminescence was measured on Tecan Plate Reader Spark with an integration time of 500 ms.

**Photocrosslinking, click chemistry and in-gel fluorescence**. MDA-MB-468 cells (2x10^5^) were grown to 80% confluence on a 12-well plate. The cells were washed with 1x HBSS (Thermo Fisher) for 5 min at 37 ^º^C. Fresh HBSS (0.5 mL) was added to each well and the cells were dosed with the indicated concentrations of different probes and incubated 30 min at 37 ^º^C. The cells were cooled to 4 ^º^C and irradiated with 365 nm light for 5 min at 3.5 mW/cm^2^ (UV Crosslinker FB-UVXL-1000, Fisher Scientific). The cells were washed 2x PBS on the plate and then lysed directly with 1% SDS in PBS. The protein concentrations were determined by BCA assay (Pierce). Then equal amount of proteins were used for click reactions with a rhodamine-N_3_ (50 μM), TCEP (1 mM), TBTA (100 μM), CuSO_4_ (1 mM) for 1 h at room temperature in the dark. The SDS concentration in the click reaction mixture was controlled to be <0.5%. 4x SDS-PAGE sample buffer (Bio-Rad) was added and the lysates were loaded onto a 4-20% TGX precast SDS-PAGE gel (Bio-Rad) for electrophoresis with in-gel fluorescence imaging. After fluorescence imaging, the gel was transferred to a nitrocellulose membrane for Western blot analyses.

**Photocrosslinking, click chemistry and streptavidin pulldown.** For western blot analysis, MDA-MB-468 cells were grown on 10-cm plates to 80% confluence. The cells were washed once with 1x HBSS (Thermo Fisher) for 5 min at 37 ^º^C. Fresh HBSS (5 mL) was added and the cells were dosed with DMSO, 0.5 or 5 µM probe **4** and incubated 30 min at 37 ^º^C (one 10-cm plates per condition). The cells were cooled to 4 ^º^C and then irradiated with 365 nm for 5 min (one plate at a time). The cells were collected mechanically, washed twice with ice-cold 1x PBS. The cell pellet was lysed directly with 1% SDS in PBS (200 µL). The protein concentrations were determined by BCA assay and then normalized for click reactions with biotin-N_3_. 0.5 mg of protein from each lysate was diluted with PBS and clicked with the biotin-azide 50 µM (+ 1 mM TCEP, 100 µM TBTA, 1 mM CuSO_4_, two reactions per condition, 400 µL per reaction 1.5 hr, rt, dark,). The click reaction mixtures for each condition were combined and 400 µL MeOH + 100 µL CHCl_3_ were added to each, shaken and centrifuged at 8,000x g for 10 min. The resulting protein solids were washed with 3 x 100 µL 1:1 MeOH/CHCl_3_. Then the protein solids were resuspended in MeOH (200 µL) by sonication (bath sonicator, 5 min). CHCl_3_ (50 µL) was added to the resuspended proteins, shaken vigorously and then centrifuged at 14,000x rpm for 10 min to pellet the solids. The organic solvent was fully removed and the solids were redissolved 1% SDS in PBS (100 µL, sonicated 30 min then left in bath overnight). The protein concentrations were determined again, normalized and diluted with PBS to 0.1% SDS. The protein solution was mixed 20 µL prewashed streptavidin-agarose beads and tumbled at room temperature for overnight. The beads were washed extensively [3x 1% SDS in PBS (100 µL) then 6x PBS (100 µL)]. The bound proteins were liberated by incubation with 1 mM biotin in 1% SDS in PBS (15 µL per condition, 5 min, 95 ^º^C). The eluted proteins were loaded onto a 4-20% gradient precast gel followed by transferring to nitrocellulose membranes for western blotting.

For proteomics analysis, MDA-MB-468 cells were grown on 15-cm plates to 80% confluence. The cells were washed with 1x HBSS (5 min, 37 ^º^C). Fresh HBSS (10 mL) was added and the cells were dosed with DMSO, 0.5 µM or 5 µM probe **4**, 1 µM or 10 µM probe **5**, 1 µM or 10 µM **6** and incubated 30 min at 37 ^º^C (two 15-cm plates per condition). The cells were cooled to 4 ^º^C and then irradiated with 365 nm for 5 min (one plate at a time) at 3.5 mW/cm^2^ (UV Crosslinker FB-UVXL-1000, Fisher Scientific). The cells were collected mechanically, washed 2x with ice-cold PBS. Then the final cell pellet was lysed with 1% SDS in PBS (1.5 mL) to obtain the whole cell lysates. The protein concentrations were determined by BCA assay (Pierce). Then 3.5 mg of proteins from each lysate was diluted with PBS and clicked with the biotin-azide 50 µM (+ 1 mM TCEP, 100 µM TBTA, 1 mM CuSO_4_, two reactions per condition, 1.5 mL per reaction 1.5 hr, rt, dark). The click reactions for each condition were combined and 3 mL MeOH + 0.75 mL CHCl_3_ were added to each, vigorously vortexed for 30s and centrifuged at 8,000x g for 10 min. The resulting protein solids were washed with 3 x 3 mL 1:1 MeOH/CHCl_3_. Then the protein solids were manually transferred to 2 mL Eppendorf vials and suspended in MeOH (1.2 mL). The mixture was bath sonicated to facilitate suspension. CHCl_3_ (300 µL) was added to the resuspended proteins, shaken vigorously and then centrifuged at 14,000x rpm for 10 min to pellet the solids. The organic solvent was fully removed and the solids were redissolved 1% SDS in PBS (1 mL, sonicated 30 min then left in bath overnight). The protein concentrations were determined again, normalized and diluted to 0.1% SDS using PBS. The solution was then mixed with 60µL prewashed streptavidin-agarose beads and tumbled overnight at room temperature. The beads were washed extensively [3x 1 mL of 1% SDS in PBS then 6x PBS (1 mL)] and the biotinylated-streptavidin bound proteins were liberated by incubation with 1 mM biotin in 1% SDS in PBS (75 µL per condition, 5 min, 95 ^º^C). The eluted proteins were separated from the streptavidin beads using Spin-X centrifuge filter tubes. The beads were washed with 100 mM NH_4_HCO_3_ (3 x 100 µL). The elution and washings were combined for proteomics analysis shown below.

**LC-MS/MS analysis**. To 300 µL each of the affinity purified proteins was added 130 µL of 20% SDS and 27.5 µL of 1M triethyl ammonium bicarbonate (TEAB) to get to 5%SDS, 50mM TEAB in 457.5 µL. The samples were reduced with dithiothreitol and alkylated with idoacetamide followed by trypsin digestion on S-Trap micro (Protifi, Inc) according to the manufacturer’s suggested protocol. After digestion overnight at 37 ^o^C, the peptides were eluted with 40 μL of 50 mM TEAB, followed by 0.2% formic acid in ddI H_2_O and 50% acetonitrile in ddI H_2_O containing 0.2% formic acid. The elutions were pooled, dried in a speedvac and redissolved in 20 μL of 5% formic acid and subjected to LC/MS/MS analysis.

Each peptide elution from above was then chromatographically separated using a Dionex RSLC UHPLC system and delivered to a Q-Exactive HF mass spectrometer (Thermo Scientific) with an EasySpray Ion source and run using a 90min LC/MS method. Xcalibur version 4.0 was used to control the system. Survey mass spectra were acquired over m/z 375−1400 at 120,000 resolution (m/z 200) and data-dependent acquisition selected the top 10 most abundant precursor ions for tandem mass spectrometry by HCD fragmentation using an isolation width of 1.2 m/z, normalized collision energy of 30, and a resolution of 30,000. Dynamic exclusion was set to auto, charge state for MS/MS +2 to +7, maximum ion time 100ms, minimum AGC target of 3 x 106 in MS1 mode and 5 x 103 in MS2 mode.

Mass spectrometry data from all samples was processed using COMET/PAWS against Uniprot Human database. Comet (v. 2016.01, rev. 3)^1^ was used to search MS2 Spectra against a January 2024 version of canonical FASTA protein database containing human uniprot sequences, and concatenated sequence-reversed entries to estimate error thresholds. Comet searches for all samples were performed with trypsin enzyme specificity with monoisotopic parent ion mass tolerance set to 1.25 Da and monoisotopic fragment ion mass tolerance set at 1.0005 Da and a variable modification of +15.9949 Da on Methionine residues.

**TMT labeling and quantification**. HEK 293T cells in 6-well plates at ~80% confluency were treated with DMSO or 1 μM cmpd 27 for 20 h. Then the cells were harvested by scraping. The cell pellets were washed 2x with ice-cold PBS. The cells were then lysed in 80 μL of lysis buffer (50 mM TrisHCl, 5 mM EDTA, 150 mM NaCl, 1 mM DTT, 0.5% Nonidet P-40, pH 8.0) supplemented with 1 mM PMSF and protease inhibitor cocktail (Roche). The lysates were then processed for TMT labeling and quantification as below.

**Proteomics Sample Processing:** A total of 50 micrograms of protein per sample was digested overnight using S-Trap microcolumns (ProtiFi) per the manufacturer suggested protocol. Fifteen micrograms of peptides per sample were labeled using a TMTpro 18-plex labeling kit (Thermo Fisher). Channels 126C-127C were DMSO treatment replicates and channels 128N-129N were replicates treated with cmpd **27** (1 μM). A quality control sample consisting of 2 microliters from each labeled sample was created and 2 micrograms of the mixture was analyzed with a 125-min LC method on an Orbitrap Eclipse Tribrid mass spectrometer (Thermo Fisher) to measure total reporter ion signal per sample.

**LC-MS:** A 35 microgram equal-reporter-ion-intensity peptide mixture of the labeled samples was separated into 20 fractions by high-pH reversed liquid chromatography (Dionex NCS-3500RS UltiMate RSLCnano UPLC; Thermo Fisher) using step elutions (17, 20, 21, 22, 23, 24, 25, 26, 27, 28, 29, 30, 31, 32, 33, 34, 35, 40, 50, and 90%) of acetonitrile. Eluted fractions were separated with 140-min low-pH reverse phase chromatography and analyzed with an Orbitrap Eclipse Tribrid (Thermo Fisher) with survey scans performed in the Orbitrap (resolution 120K, m/z 375-1500, AGC target 400K, and max ion inject time 246ms), and data-dependent MS2 scans in the linear ion trap using CID (1 Da isolation window, NCE 35%, max ion inject time 75ms, AGC target of 20K, and Rapid mode). Reporter ion detection was performed in the Orbitrap using MS3 scans (resolution 50K, NCE 55%, max ion inject time 86ms, AGC target 100K, and scan range 100-150) following synchronous precursor isolation (ten 2-Da notches) in the linear ion trap, and HCD in the instrument’s ion-routing multipole. Cycle time was 2 sec with 10ppm 45-sec dynamic exclusion enabled.

**Data processing:**

The binary instrument files were converted to compressed text files using MSConvert, part of the ProteoWizard toolkit (https://proteowizard.sourceforge.io/). PAW pipeline Python scripts (<https://github.com/pwilmart/PAW_pipeline>)^2^ extracted TMTpro reporter ion peak heights and fragment ion spectra in MS2 format. The Comet search engine (v. 2016.03)^1^ used 1.25 Da monoisotopic peptide mass tolerance, 1.0005 Da monoisotopic fragment ion tolerance, semi tryptic cleavage with up to 3 missed cleavages, variable oxidation of methionine and proline residues, static alkylation of cysteines, and static modifications for TMTpro labels (at peptide N-termini and lysine residues). The FASTA file was the UniProt Homo sapiens reference proteome (UP000005640, taxon ID 9606, one protein per gene, 20,598 protein sequences downloaded April 2024) appended with 175 common contaminant sequences, and concatenated sequence-reversed decoy entries.

Approximately 542K MS2/MS3 spectra were acquired from the 20 fractions, of which 38.1% were matched to peptide sequences at a 1% false discovery rate determined using the target/decoy method. The resulting 206K peptide sequences mapped to 6,758 proteins (2 peptides per protein) using basic and extended parsimony rules. After excluding decoys, common contaminants, and proteins without usable reporter ion signals, a final set of 6,608 quantifiable proteins was obtained. Normalization of reporter ion intensities was performed using the trimmed mean of M-values (TMM) method implemented in the Bioconductor package edgeR.^3^ Data quality was assessed by examining box plots of TMM-normalized protein intensities, sample-to-sample correlation plots, and MDS clustering plots prior to further differential analysis of protein abundance between biological groups using the exact test in edgeR with Benjamini-Hochberg multiple testing corrections.

**CeTSA**. MDA-MB-468 cells (80% confluence, 10-cm plate) were incubated with DMSO or **8** (5 µM) for 3 hours at 37 ^º^C. The cells were collected mechanically and washed with ice-cold PBS (2x 5 mL) and centrifuged at 200x g for 5 min. The final cell pellets were resuspended in 80 µL of PBS and 10 µL of the cell suspension was aliquoted into PCR tubes. The samples were heated at the indicated temperatures (30-100 ^º^C) for 3 min followed by 25 ^º^C for 3 min in a PCR thermal cycler. Then 20 µL HEPES buffered saline (20 mM HEPES, 250 mM NaCl, pH 8.0) supplemented with protease inhibitor cocktail (Roche) and 1 mM PMSF was added to the cell suspension, which was then subjected to three cycles of freeze/thaw in liquid nitrogen (each freezing time was extended to 10 min in liquid N_2_). The lysates were centrifuged at 14,000 rpm for 15 min at 4 ^º^C and supernatant was separated from the insoluble components. To the remaining insoluble components, 1% SDS in PBS (45 µL) was added and the contents were briefly sonicated with a probe sonicator. The lysates were then centrifuged at 14,000x rpm for 10 min. The samples were mixed with 2x SDS-PAGE buffer with 5% beta-mercaptoethanol (Bio-Rad) and heated at 95 ^º^C for 5 min, and then separated on a 4-20% gel by SDS-PAGE and transferred to a nitrocellulose membrane for analysis by Western blot.

**sh*SYVN1* knockdown**. HEK293T cells were transfected with lentiviral expression plasmids (sh*CTRL* and sh*SYVN1* (Sigma, TGAATGCTTAATCCCGGGAAA)) along with packaging vectors using the calcium-phosphate method. The supernatants containing lentiviral particles were collected, passed through a 0.45 μM filter, and stored at −80 °C prior to use. For lentiviral transduction, HEK293T were transduced with sh*CTRL* or sh*SYVN1* lentiviruses for 24 h. A second round of transduction using the same lentiviruses was conducted for another 24 h. The transduced cells were maintained in puromycin (0.6 µg/mL)-containing media selection.

**BRD4 degradation assay**. The cells were plated into 12-well plate and allowed to attach over 3 hr in media. The cells were then treated with DMSO or different concentrations of **27** for the 2 or 20 hr at 37 ^º^C. In certain experiments, MG132 or JQ1 or compound **8** were added the same time. The cells were then collected by scraping, washed with ice-cold PBS. The cell pellet was lysed lysis buffer (50 mM TrisHCl, 5 mM EDTA, 150 mM NaCl, 1 mM DTT, 0.5% Nonidet P-40, pH 8.0) supplemented with protease inhibitor cocktail (Roche) and 1 mM PMSF. The protein concentration was determined using Dye Reagent Concentrate (Bio-Rad). Then equal amounts of proteins were loaded onto 4-20% precast SDS-PAGE for electrophoresis followed by electrotransfer and Western blot analyses.

**Immunofluorescence**. HEK293T cells transduced with sh*CTRL* or sh*SYVN1* were grown on 2% Matrigel (Corning) coated coverslips at 37 ºC. The cells were washed twice with 0.5 mL DPBS (Thermo Fisher). The cells were fixed with 4% PFA (in serum free DMEM) and permeabilized with 0.1% triton-X-100. The cells were blocked in 3% BSA in PBS for 1 hr and then incubated with anti-SYVN1 (Bethyl laboratories A302-946A,1:500) in 3% BSA overnight at 4 ºC. The cells were washed and then incubated with anti-rabbit AlexaFluor488 (1:1000 in 3% BSA, dark, rt, 1 hr). The nuclei were stained with DAPI (1:2000 dilution, 10 min, rt). The cells were washed with PBS and the coverslips were mounted onto glass slides. The coverslips were dried overnight prior to confocal image acquisition using Zeiss LSM980.

**Chemistry.** All ^1^H and ^13^C NMR spectra were obtained in a Bruker Avance-Neo 400 MHz spectrometer in CDCl_3_ or DMSO-*d*_6_ using the chemical shifts of the residual CHCl_3_ (*δ* 7.26) or DMSO (*δ* 2.50) as the reference. Chemical shifts (*δ*) are reported in parts per million (ppm), and the signal multiplicity is reported as brs (broad singlet), d (doublet), dd (doublet of doublet), td (triplet of doublet), m (multiplet), q (quartet), s (singlet), and t (triplet). Coupling constants (*J* values) are given in Hz. Purification by flash chromatography was performed using 230-400 mesh silica gel (EMD). Reaction progress was followed by thin-layer chromatography (TLC) on silica gel plates (EMD). Percent yield of final purified compounds is reported. Melting points were determined in capillary tubes using Mel-Temp and are uncorrected. The mass spectra of the small molecules were obtained from Advion PlateExpress by electrospray ionization in both positive and negative modes.

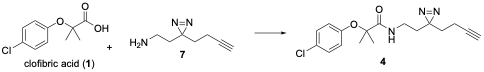

***N*-(2-(3-(but-3-yn-1-yl)-3*H*-diazirin-3-yl)ethyl)-2-(4-chlorophenoxy)-2-methylpropanamide** (**4**).

Clorfibric acid (10 mg, 0.05mmol), EDCI (13 mg, 0.06 mmol) and Et_3_N (0.1 mmol, 14 µL) were dissolved in 0.5 mL dry DMF. Compound **7** (5 mg, 0.04 mmol) was added and the reaction mixture was stirred at rt for 3 hours in the dark. Saturated sodium bicarbonate (aq, 20 mL) was added and the reaction mixture was extracted with DCM (3 x 10 mL). The combined organic extracts were washed with brine (1x 5 mL), dried over Na_2_SO_4_ and concentrated. The crude isolate was purified by flash chromatography eluting with ether to afford **4** as an oily yellow solid (3.9 mg, 45%). ^1^H NMR (400 MHz, CDCl_3_) *δ* 7.25 (d, *J* = 9.0 Hz, 2H), 6.88 (d, *J* = 8.9 Hz, 2H), 6.75 (brs, 1H), 3.21 (q, *J* = 6.8 Hz, 2H), 2.08 – 1.90 (m, 3H), 1.71 (t, *J* = 6.8 Hz, 2H), 1.64 (t, *J* = 7.1 Hz, 2H), 1.50 (s, 6H). ^13^C NMR (101 MHz, CDCl_3_) *δ* 174.46, 152.75, 129.30, 128.67, 122.76, 81.88, 69.41, 34.34, 32.56, 32.04, 29.71, 26.74, 24.90, 13.19. APCI Calcd for [C_17_H_21_ClN_3_O_2_]^+^ 334.1; found 334.2.

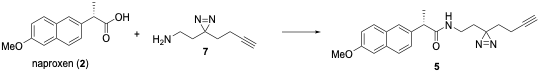

***N*-(2-(3-(but-3-yn-1-yl)-3*H*-diazirin-3-yl)ethyl)-2-(6-methoxynaphthalen-2-yl)propenamide** (**5**).

Naproxen (6.1 mg, 0.026mmol) and EDCI (4.1mg, 0.026 mmol) were dissolved in 0.6 mL THF. Compound **7** (3 mg, 0.022 mmol) was added and the reaction mixture was stirred at rt for 3 hours in the dark. Saturated sodium bicarbonate (aq, 10 mL) was added and the reaction mixture was extracted with DCM (3 x 5 mL). The combined organic extracts were washed with brine (1x 5 mL), dried over Na_2_SO_4_ and concentrated. The crude was purified by flash chromatography eluting with DCM to afford **5** as a white solid (4.6 mg, 84%). Mp 139-142 ^o^C.

^1^H NMR (400 MHz, CDCl_3_) *δ* 7.73 (d, *J* = 8.4 Hz, 1H), 7.72 (d, *J* = 8.8 Hz, 1H), 7.68 (s,1H), 7.39 (dd, *J* = 8.4, 2.0 Hz, 1H), 7.16 (dd, *J* = 8.8, 2.4 Hz, 1H), 7.12 (d, *J* = 2.4 Hz, 1H), 5.46 (brs, 1H), 3.92 (s, 3H), 3.69 (q, *J* = 7.2 Hz, 1H), 3.14 – 2.78 (m, 2H), 1.91 – 1.84 (m, 3H), 1.61 (d, *J* = 7.3 Hz, 3H), 1.57 (t, *J* = 6.4 Hz, 2H), 1.51 (d, *J* = 7.1 Hz, 1H). ^13^C NMR (101 MHz, CDCl_3_) *δ* 174.49, 157.94, 136.34, 133.95, 129.37, 129.14, 127.79, 126.~~45~~30, 126.16, 119.40, 105.80, 82.76, 69.40, 55.49, 47.25, 34.64, 32.63, 32.16, 26.83, 18.43, 13.25. APCI+ Calcd for [C_21_H_23_N_3_O_2_]^+^ [M+H]^+^350.2; found 350.~~4~~2

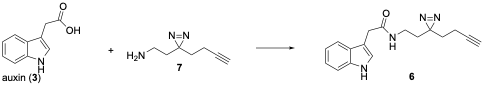

***N*-(2-(3-(but-3-yn-1-yl)-3*H*-diazirin-3-yl)ethyl)-2-(1*H*-indol-3-yl)acetamide** (**6**).

 Auxin (6.6 mg, 0.053mmol) and EDCI (12.1mg, 0.063 mmol) were dissolved in 0.5 mL THF. Et_3_N (0.06 mmol, 6 µL) was added. Compound **7** (5 mg, 0.04 mmol) was dissolved in 0.25 mL THF and added to the reaction mixture over 1hr at rt. The reaction mixture was stirred at rt for 1 hr then 0.25 mL DMF was added to improve solubility and stirred an addition 30 min. Saturated sodium bicarbonate (aq, 15 mL) was added and the reaction mixture was extracted with DCM (3 x 10 mL). The combined organic extracts were washed with brine (1x 5 mL), dried over Na_2_SO_4_ and concentrated. The crude was purified by filtration over a silica plug and the silica plug was eluted with ether to afford **6** as an oily brown solid (4.4 mg, 60%). ^1^H NMR (400 MHz, CDCl_3_) *δ* 8.29 (brs, 1H), 7.61 (d, *J* = 8.0 Hz, 1H), 7.44 (d, *J* = 8.0 Hz, 1H), 7.25 – 7.10 (m, 3H), 5.79 (brs, 1H), 3.75 (s, 2H), 3.05 (q, *J* = 6.6 Hz, 2H), 1.89 – 1.77 (m, 3H), 1.54 (t, *J* = 6.7 Hz, 2H), 1.49 (t, *J* = 7.4 Hz, 2H). ^13^C NMR (101 MHz, CDCl_3_) *δ* 171.64, 136.56, 127.16, 123.95, 122.89, 120.33, 118.88, 111.57, 109.11, 82.74, 69.32, 34.50, 33.52, 32.70, 32.08, 26.79, 13.19. ESI Calcd for [C_17_H_1~~9~~7_N_4_O]^+^ [M-H]^-^ 293.1 found 292.9.

 
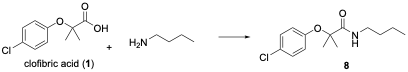

***N*-butyl-2-(4-chlorophenoxy)-2-methylpropanamide (8).**

Clofibric acid (25 mg, 0.117 mmol), butylamine (10.2 mg, 0.140 mmol, 15µL), EDCI (26 mg, 0.140) and Et_3_N (14 mg, 0.140 mmol, 20 µL) were dissolved in DCM (1 mL). The reaction mixture was stirred at rt for 1.5 hr. Aqueous NaHCO_3_ (sat, 5 mL) was added and stirred for 5 min at rt. The aqueous mixture was diluted with water (to 10 mL) and extracted with DCM (3 x 3 mL). The combined organic extracts were dried over Na_2_SO_4_ and concentrated. The crude mixture was then filtered through a silica plug in Et_2_O. The filtrate was concentrated to afford the product, **8**, as a clear, colorless, viscous oil (15.4 mg, 50% yield). ^1^H NMR (400 MHz, CDCl_3_) *δ* 7.23 (d, *J* = 9.0 Hz, 2H), 6.84 (d, *J* = 8.9 Hz, 2H), 3.29 (td, *J* = 7.1, 5.9 Hz, 2H), 1.51-1.43 (m, 8H), 1.32 (quintet, *J* = 6.9 Hz, 2H), 0.91 (t, *J* = 7.3 Hz, 3H). ^13^C NMR (101 MHz, CDCl_3_) *δ* 174.27, 152.91, 129.23, 128.41, 122.46, 81.89, 39.10, 31.58, 25.05, 20.05, 13.72. ESI+ Calcd for [C_14_H_21_ClNO_2_]^+^ 270.1; found 270.1

**2-(4-Chlorophenoxy)-*N*-(9-(3-(5,5-difluoro-7-(1*H*-pyrrol-2-yl)-5*H*-4*λ*^4^,5 *λ*^4^-dipyrrolo[1,2-*c*:2',1'-*f*][1,3,2]diazaborinin-3-yl)propanamido)nonyl)-2-methylpropanamide (9):**

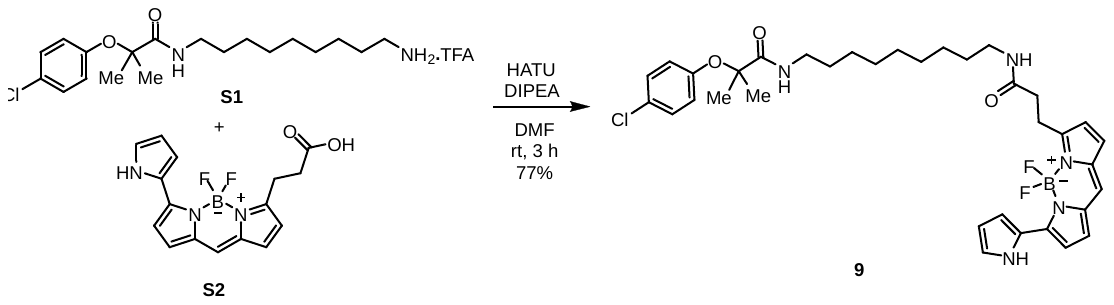

Acid **S2** (21 mg, 0.06 mmol, 1.0 equivalent), DIPEA (33 µL, 3.0 equivalents) and HATU (36 mg, 1.5 equivalents) were added to a stirred solution of amine **S1** (30 mg, 1.0 equivalent) in DMF (1.5 mL) at room temperature. The reaction mixture was covered with aluminum foil and stirred for 3 h at room temperature. The mixture was then quenched with sat aq NH_4_Cl (7 mL) and extracted with EtOAc (2 X 15 mL). The combined organic layers were dried over sodium sulfate. The solution was filtered, concentrated and the residue was purified by silica gel column chromatography (0-2% MeOH in DCM) to give probe **9** in 77% (33 mg) yield as a dark bluish fluorescent oil.

^1^H NMR (400 MHz, CDCl_3_) *δ* 10.39 (brs, 1H), 7.23 (d, *J* = 6.8 Hz, 2H), 7.19 – 7.15 (m, 1H), 7.04 (d, *J* = 4.4 Hz, 1H), 7.01 – 6.97 (m, 2H), 6.87 (d, *J* = 4.4 Hz, 1H), 6.86 – 6.82 (m, 3H), 6.62 (brs, 1H), 6.40 – 6.36 (m, 1H), 6.30 (d, *J* = 4.0 Hz, 1H), 5.60 (brs, 1H), 3.35 – 3.16 (m, 6H), 2.64 (t, *J* = 7.6 Hz, 2H), 1.50 – 1.46 (m, 10H), 1.43 – 1.38 (m, 2H), 1.28 – 1.17 (m, 8H). ^13^C NMR (101 MHz, CDCl_3_) *δ* 174.39, 171.66, 155.53, 153.04, 150.59, 137.44, 133.65, 131.70, 129.37, 128.49, 126.56, 125.81, 123.81, 123.48, 122.51, 120.44, 117.89, 117.15, 111.66, 81.97, 39.64, 39.48, 36.15, 29.59, 29.44, 29.24, 29.23, 26.93, 26.85, 25.19, 24.91. ESI-MS: Calcd for C_35_H_44_BClF_2_N_5_O_3_ [M+H]^+^ 666.3, Found [M+H]^+^ 666.8.

**Synthesis of PROTAC 10:**

***Tert*-butyl (2-aminoethyl)carbamate S4:**

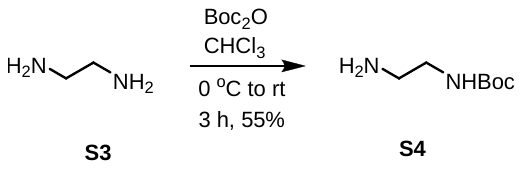

A solution of Boc_2_O (1 g, 4.6 mmol, 1 equiv) in CHCl_3_ (30 mL) was added dropwise to a solution of the diamine **S3** (2.45 mL, 36.6 mmol, 8 equiv) in CHCl_3_ (7 mL) at 0 °C over a 30 min period. The reaction mixture was then stirred at room temperature for 3 h. The mixture was filtered, and the filtrate was concentrated under reduced pressure. The residue was azeotroped with toluene (2 X 2 mL). The crude residue was dissolved in EtOAc (30 mL) and washed with brine (10 mL). The organic solution was dried over Na_2_SO_4_ and concentrated under reduced pressure to give carbamate **S4** in 55% (400 mg) as a colorless oil. ^1^H NMR (400 MHz, CDCl_3_) *δ* 4.92 (brs, 1H), 3.20 – 3.151 (m, 2H), 2.83 – 2.76 (m, 2H), 1.43 (s, 9H).

**(*S*)-*N*-(2-Aminoethyl)-2-(4-(4-chlorophenyl)-2,3,9-trimethyl-6H-thieno[3,2-f][1,2,4]triazolo[4,3-a][1,4]diazepin-6-yl)acetamide (S6):**

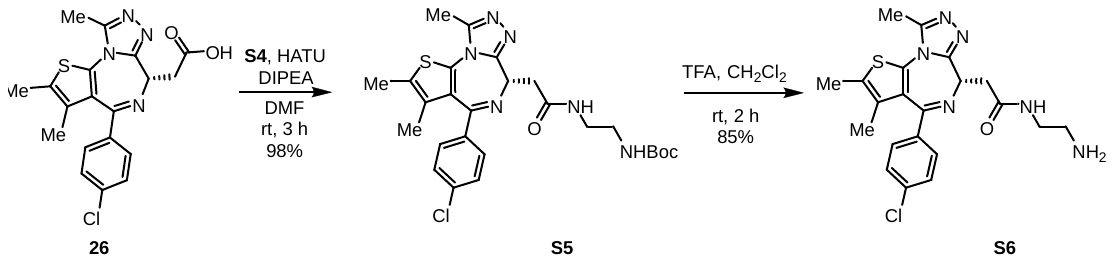

Acid **26** (100 mg, 0.25 mmol, 1.0 equivalents), DIPEA (174 µL, 4.0 equivalents) and HATU (190 mg, 2.0 equivalents) were added to a stirred solution of carbamate **S4** (48 mg, 0.3 mmol, 1.2 equivalents) in DMF (1.3 mL) at room temperature. The mixture was stirred for 3 h at room temperature. The mixture was then diluted with water (7 mL) and extracted with EtOAc (2 X 10 mL). The combined organic layers were washed with brine (5 mL), and the solution was dried over sodium sulphate. The solution was filtered, concentrated and the residue was purified by silica gel column chromatography (0-6% MeOH/DCM) to give compound **S5** in 98% (134 mg) as a colorless solid.

^1^H NMR (400 MHz, CDCl_3_) *δ* 7.43 (d, *J* = 8.0 Hz, 2H), 7.35 (d, *J* = 8.0 Hz, 2H), 7.14 (brs, 1H), 5.21 (brs, 1H), 4.69 (t, *J* = 8.0 Hz, 1H), 3.53 (dd, *J* = 12.0, 4.0 Hz, 1H), 3.42 – 3.34 (m, 3H), 3.27 – 3.25 (m, 2H), 2.69 (s, 3H), 2.41 (s, 3H), 1.68 (s, 3H), 1.42 (s, 9H). ESI-MS: Calcd for C_27_H_32_N_5_O_3_SCl [M+H]^+^ 543.2, Found [M+H]^+^ 543.2.

Compound **S5** (134 mg, 0.25 mmol) was dissolved in DCM (1.5 mL)., Then TFA (150 µL) (10:1 DCM:TFA) was added at room temperature. The reaction mixture was stirred for 2 hours at the same temperature. Afterward, the volatiles were evaporated, and the crude reaction mixture was dissolved in 10% MeOH in DCM (5 mL) and quenched with NaHCO_3_ (7 mL). The mixture was then extracted with 10% MeOH in DCM (2 × 10 mL), and the combined organic layers were dried over Na_2_SO_4_. The solution was filtered, concentrated under reduced pressure, and amine **S6** was obtained as a colorless oil in 85% (93 mg). This compound was used in subsequent reactions without further purification.

^1^H NMR (400 MHz, CDCl_3_) *δ* 7.41 (d, *J* = 8.4 Hz, 2H), 7.33 – 7.30 (m, 2H), 4.66 (dd, *J* = 8.0, 6.0 Hz, 1H), 3.60 – 3.50 (m, 3H), 3.41 – 3.34 (m, 3H), 2.97 (brs, 1H), 2.66 (s, 3H), 2.40 (s, 3H), 1.68 (s, 3H). ESI-MS: Calcd for C_21_H_24_N_6_OSCl [M+H]^+^ 443.1, Found [M+H]^+^ 443.2.

**(*S*)-2-(4-chlorophenoxy)-*N*-(2-(2-(4-(4-chlorophenyl)-2,3,9-trimethyl-6*H*-thieno[3,2-*f*][1,2,4]triazolo[4,3-*a*][1,4]diazepin-6-yl)acetamido)ethyl)-2-methylpropanamide (10):**

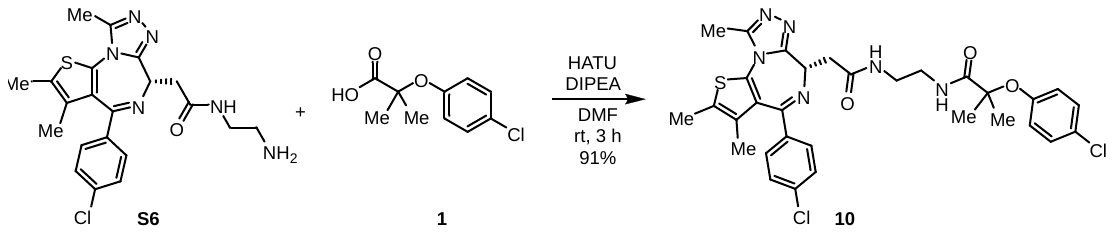

Amine **S6** (28 mg, 0.06 mmol, 1.0 equivalents) was dissolved in DMF (1 mL), followed by the addition of clofibric acid (**1**, 16.5 mg, 1.2 equivalents), DIPEA (44 µL, 4.0 equivalents), and HATU (48 mg, 2.0 equivalents) to the mixture at room temperature. The mixture was stirred for 2 hours at room temperature. The mixture was then diluted with water (5 mL) and extracted with EtOAc (2 × 10 mL). The combined organic layers were washed with brine (5 mL), and the solution was dried over sodium sulfate. The solution was filtered, concentrated, and the residue was purified by silica gel column chromatography (0-4% MeOH/DCM) to give compound **10** in 91% yield (39 mg) as a colorless solid.

^1^H NMR (400 MHz, CDCl_3_) *δ* 7.56 (brs, 1H), 7.39 (d, *J* = 8.0 Hz , 2H), 7.32 – 7.30 (m, 2H), 7.17 – 7.13 (m, 2H), 6.84 – 6.79 (m, 2H), 4.61 (t, *J* = 6.8, Hz, 1H), 3.49 – 3.24 (m, 6H), 2.63 (s, 3H), 2.40 (s, 3H), 1.66 (s, 3H), 1.48 (s, 3H), 1.47 (s, 3H). ^13^C NMR (101 MHz, Chloroform-*d*) *δ* 175.15, 171.33, 164.24, 155.65, 153.35, 150.06, 137.14, 136.41, 132.16, 131.26, 131.14, 130.58, 130.07, 129.28, 128.89, 128.11, 122.28, 81.71, 54.28, 39.85, 39.50, 38.87, 25.49, 25.31, 24.83, 14.51, 13.25, 11.86. ESI-MS: Calcd for C_31_H_33_N_6_O_3_SCl_2_ [M+H]^+^ 639.2, Found [M+H]^+^ 639.3. M.P: 89-91 ^o^C.

**Synthesis of PROTAC 11:**

***Tert*-butyl (3-aminopropyl)carbamate (S8):**

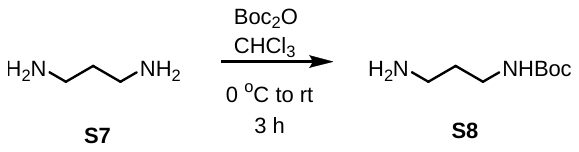

A solution of Boc_2_O (2 g, 9.2 mmol, 1 equiv) in CHCl_3_ (30 mL) was added dropwise to a solution of the diamine **S7** (6.2 mL, 73.3 mmol, 8 equiv) in CHCl_3_ (13 mL) at 0 °C over a 30 min period. The reaction mixture was then stirred at room temperature for 3 h. The mixture was filtered, and the filtrate was concentrated under reduced pressure. The residue was azeotroped with toluene (2 X 2 mL). The crude residue was dissolved in EtOAc (30 mL) and washed with brine (10 mL). The organic solution was dried over Na_2_SO_4_ and concentrated under reduced pressure to give carbamate **S8** as a light yellowish oil (1.5 g, 99%). ^1^H NMR (400 MHz, CDCl_3_) *δ* 4.93 (brs, 1H), 3.19 (q, *J* = 6.7 Hz, 2H), 2.74 (t, *J* = 6.8 Hz, 2H), 1.58 (quintet, *J* = 6.8 Hz, 2H), 1.42 (s, 9H), 1.21 (brs, 2H).

**(*S*)-*N*-(3-Aminopropyl)-2-(4-(4-chlorophenyl)-2,3,9-trimethyl-6*H*-thieno[3,2-*f*][1,2,4]triazolo[4,3-*a*][1,4]diazepin-6-yl)acetamide (S10):**

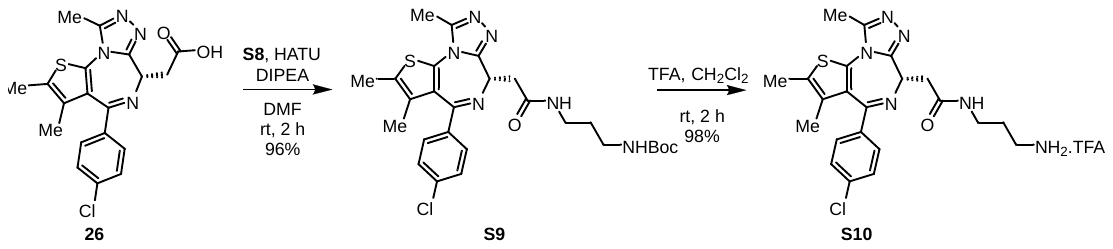

Acid **26** (100 mg, 0.25 mmol, 1.0 equivalents), DIPEA (217 µL, 5.0 equivalents) and HATU (190 mg, 2.0 equivalents) were added to a stirred solution of *tert*-butyl (3-aminopropyl)carbamate (**S8**, 52 mg, 0.3 mmol, 1.2 equivalents) in DMF (1.3 mL) at room temperature. The mixture was stirred for 2 h at room temperature. The mixture was then diluted with water (7 mL) and extracted with EtOAc (2 X 10 mL). The combined organic layers were washed with brine (5 mL), and the solution was dried over anhydrous sodium sulfate. The solution was filtered, concentrated and the residue was purified by silica gel column chromatography (0-6% MeOH/DCM) to give compound **S9** as a yellow solid (133 mg, 96%).

^1^H NMR (400 MHz, CDCl_3_) *δ* 7.40 (d, *J* = 8.0 Hz, 2H), 7.32 (d, *J* = 8.0 Hz, 2H), 6.92 (brs, 1H), 5.14 (brs, 1H), 4.64 (t, *J* = 6.9 Hz, 1H), 3.55 (dd, *J* = 12.0, 4.0 Hz, 1H), 3.42 – 3.24 (m, 3H), 3.15 (q, *J* = 4.0 Hz, 2H), 2.67 (s, 3H), 2.40 (s, 3H), 1.60-1.70 (m, 5H), 1.43 (s, 9H). ESI-MS: Calcd for C_27_H_34_N_6_O_3_SCl [M+H]^+^ 557.2, Found [M+H]^+^ 557.3.

Compound **S9** (120 mg, 0.2 mmol) was dissolved in DCM (1.5 mL). Then TFA (150 µL) (10:1 DCM:TFA) was added at room temperature. The reaction mixture was stirred for 2 hours at the same temperature. Afterward, the volatiles were then evaporated, and DCM (3 × 2 mL) was added to the residue. The mixture was re-evaporated three times. Finally, the residue was washed with hexanes and concentrated under reduced pressure, producing compound **S10** as a dark yellowish viscous oil (120 mg, 98%). This compound was used in subsequent reactions without further purification.

^1^H NMR (400 MHz, CDCl_3_) *δ* 7.74 (brs, 1H), 7.40 (d, *J* = 8.4 Hz, 2H), 7.31 (dd, *J* = 8.4, 3.2 Hz, 2H), 4.65 (t, *J* = 7.2 Hz, 1H), 4.01 (brs, 2H), 3.69 – 3.53 (m, 1H), 3.52 – 3.41 (m, 1H), 3.36 – 3.25 (m, 2H), 2.89 (brs, 2H), 2.65 (s, 3H), 2.39 (s, 3H), 1.77 (brs, 2H), 1.66 (s, 3H). ESI-MS: Calcd for C_22_H_26_N_6_OSCl [M+H]^+^ 457.1, Found [M+H]^+^ 457.2.

**(*S*)-2-(4-Chlorophenoxy)-*N*-(3-(2-(4-(4-chlorophenyl)-2,3,9-trimethyl-6*H*-thieno[3,2-*f*][1,2,4]triazolo[4,3-*a*][1,4]diazepin-6-yl)acetamido)propyl)-2-methylpropanamide (11):**

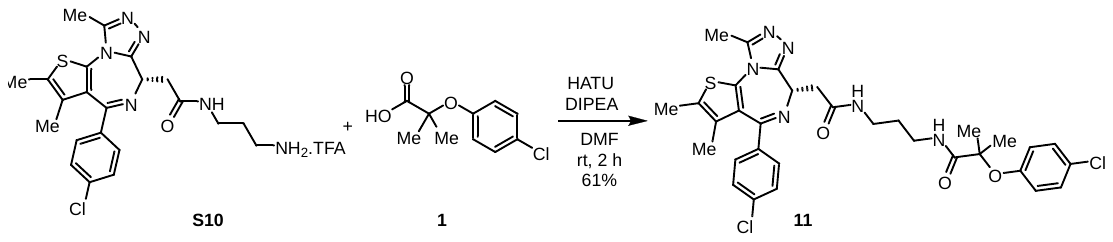

Amine **S10** (45 mg, 0.08 mmol, 1.0 equivalents) was dissolved in DMF (1 mL), followed by the addition of clofibric acid (**1**, 20.5 mg, 1.2 equivalents), DIPEA (55 µL, 4.0 equivalents), and HATU (60 mg, 2.0 equivalents) to the mixture at room temperature. The mixture was stirred for 2 hours at room temperature. The mixture was then diluted with water (7 mL) and extracted with EtOAc (2 × 10 mL). The combined organic layers were washed with brine (5 mL), and the solution was dried over sodium sulfate. The solution was filtered, concentrated, and the residue was purified by silica gel column chromatography (0-4% MeOH/DCM) to give compound **11** in 61% yield (32 mg) as a colorless oil.

^1^H NMR (400 MHz, CDCl_3_) *δ* 7.34-7.40 (m, 3H), 7.30 (d, *J* = 8.4 Hz, 2H), 7.17 (d, *J* = 8.8 Hz, 2H), 7.04 (brs, 1H), 6.84 (d, *J* = 8.8 Hz, 2H), 4.61 (dd, *J* = 7.2, 6.8 Hz, 1H), 3.52 (dd, *J* = 14.4, 7.6 Hz, 1H), 3.40 – 3.24 (m, 5H), 2.64 (s, 3H), 2.40 (s, 3H), 1.69-1.67 (m, 2H), 1.65 (s, 3H), 1.48 (s, 6H). ^13^C NMR (101 MHz, CDCl_3_) *δ* 174.81, 171.13, 164.05, 155.79, 153.24, 150.06, 136.97, 136.65, 132.18, 131.05, 131.02, 130.63, 129.94, 129.29, 128.86, 128.26, 122.49, 81.82, 54.51, 39.35, 36.44, 36.18, 29.62, 25.31, 25.08, 14.49, 13.23, 11.93. ESI-MS: Calcd for C_32_H_35_N_6_O_3_SCl_2_ [M+H]^+^ 653.2, Found [M+H]^+^ 653.3.

**Synthesis of PROTAC 12:**

***Tert*-butyl (4-aminobutyl)carbamate (S12):**

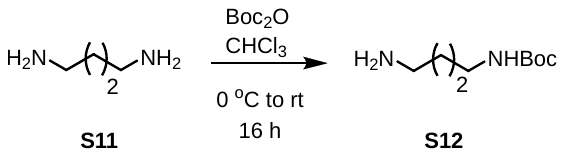

A solution of Boc_2_O (500 mg, 2.29 mmol, 1 equivalent) in CHCl_3_ (12 mL) was added dropwise to a solution of the diamine **S11** (1.6 g, 18.3 mmol, 8 equiv) in CHCl_3_ (18 mL) at 0 °C over a 30-min period. The reaction mixture was then stirred at room temperature for 16 h. The mixture was filtered, and the filtrate was concentrated under reduced pressure. The crude residue was dissolved in DCM (20 mL) and washed with brine (7 mL). The organic solution was dried over Na_2_SO_4_ and concentrated under reduced pressure to give carbamate **S12** in 95% yield (412 mg) as a colorless oil.

^1^H NMR (400 MHz, CDCl_3_) *δ* 4.67 (brs, 1H), 3.11 (q, *J* = 6.0 Hz, 2H), 2.70 (t, *J* = 6.2 Hz, 2H), 1.59 – 1.40 (m, 13H), 1.28 (brs, 2H).

**(*S*)-*N*-(4-aminobutyl)-2-(4-(4-chlorophenyl)-2,3,9-trimethyl-6*H*-thieno[3,2-*f*][1,2,4]triazolo[4,3-*a*][1,4]diazepin-6-yl)acetamide (S14):**

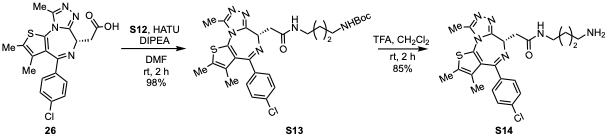

Acid **26** (100 mg, 0.25 mmol, 1.0 equivalents), DIPEA (174 µL, 4.0 equivalents) and HATU (190 mg, 2.0 equivalents) were added to a stirred solution of carbamate **S12** (56 mg, 0.3 mmol, 1.2 equivalents) in DMF (1.3 mL) at room temperature. The mixture was stirred for 2 h at room temperature. The mixture was then diluted with water (7 mL) and extracted with EtOAc (2 X 10 mL). The combined organic layers were washed with brine (5 mL), and the solution was dried over sodium sulphate. The solution was filtered, concentrated and the residue was purified by silica gel column chromatography (0-5% MeOH/DCM) to give compound **S13** in 98% (140 mg) as a colorless solid.

^1^H NMR (400 MHz, CDCl_3_) *δ* 7.42 (d, *J* = 8.8 Hz, 1H), 7.34 (d, *J* = 8.4 Hz, 1H), 6.72 (s, 1H), 4.71 (s, 1H), 4.68 – 4.61 (m, 1H), 3.54 (dd, *J* = 16.0, 8.0 Hz, 1H), 3.36 – 3.24 (m, 3H), 3.10 (brs, 2H), 2.68 (s, 3H), 2.40 (s, 3H), 1.67 (s, 3H), 1.55 – 1.47 (m, 4H), 1.43 (s, 9H). ESI-MS: Calcd for C_28_H_36_N_6_O_3_SCl [M+H]^+^ 571.2, Found [M+H]^+^ 571.1.

Compound **S13** (140 mg, 0.24 mmol) was dissolved in DCM (1.8 mL). Then TFA (180 µL) (10:1 DCM:TFA) was added at room temperature. The reaction mixture was stirred for 2 hours at the same temperature. Afterward, the volatiles were evaporated, and the crude reaction mixture was dissolved in 10% MeOH in DCM (5 mL) and quenched with NaHCO_3_ (7 mL). The mixture was then extracted with 10% MeOH in DCM (2 × 10 mL), and the combined organic layers were dried over Na_2_SO_4_. The solution was filtered, concentrated under reduced pressure, and amine **S14** was obtained as a light-yellow solid in 85% (98 mg). This compound was used in subsequent reactions without further purification.

^1^H NMR (400 MHz, CDCl_3_) *δ* 7.71 (brs, 1H), 7.41 (d, *J* = 8.0 Hz, 2H), 7.33 (d, *J* = 8.0 Hz, 2H), 4.72 – 4.59 (m, 1H), 3.71 – 3.47 (m, 2H), 3.37 – 3.25 (m, 2H), 3.14 – 3.00 (m, 2H), 2.67 (d, *J* = 4.0 Hz, 3H), 2.40 (s, 3H), 1.84 - 1.59 (m, 7H). (note: one of the methyls show 2 peaks) ESI-MS: Calcd for C_23_H_28_N_6_OSCl [M+H]^+^ 471.0, Found [M+H]^+^ 471.2.

**(*S*)-2-(4-chlorophenoxy)-*N*-(4-(2-(4-(4-chlorophenyl)-2,3,9-trimethyl-6*H*-thieno[3,2-*f*][1,2,4]triazolo[4,3-*a*][1,4]diazepin-6-yl)acetamido)butyl)-2-methylpropanamide (12):**

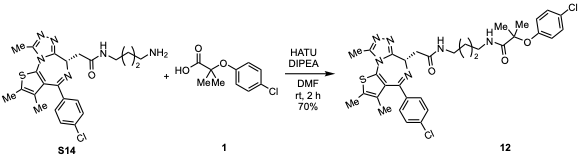

Amine **S14** (31 mg, 0.06 mmol, 1.0 equivalents) was dissolved in DMF (1 mL), followed by the addition of clofibric acid (**1**, 17 mg, 1.2 equivalents), DIPEA (46 µL, 4.0 equivalents), and HATU (50 mg, 2.0 equivalents) to the mixture at room temperature. The mixture was stirred for 2 hours at room temperature. The mixture was then diluted with water (5 mL) and extracted with EtOAc (2 X 10 mL). The combined organic layers were washed with brine (5 mL), and the solution was dried over sodium sulfate. The solution was filtered, concentrated, and the residue was purified by silica gel column chromatography (0-4% MeOH/DCM) to give compound **12** in 70% yield (30 mg) as a colorless solid.

^1^H NMR (400 MHz, CDCl_3_) *δ* 7.40 (d, *J* = 8.0 Hz, 2H), 7.33 (d, *J* = 8.0 Hz, 2H), 7.24 – 7.19 (m, 2H), 6.88 – 6.82 (m, 3H), 6.72 (s, 1H), 4.63 (t, *J* = 8.0 Hz, 1H), 3.56 (dd, *J* = 14.4, 8.0 Hz, 1H), 3.37 – 3.25 (m, 5H), 2.65 (s, 3H), 2.40 (s, 3H), 1.66 (s, 3H), 1.59 – 1.51 (m, 4H), 1.49 (s, 3H), 1.48 (s, 3H). ^13^C NMR (101 MHz, CDCl_3_) *δ* 174.56, 170.53, 164.28, 155.62, 153.04, 150.08, 137.16, 136.44, 132.22, 131.24, 131.14, 130.56, 130.03, 129.36, 128.91, 128.52, 122.74, 81.98, 54.57, 39.40, 39.26, 39.10, 27.01, 26.89, 25.17, 14.51, 13.24, 11.91. ESI-MS: Calcd for C_33_H_37_N_6_O_3_SCl_2_ [M+H]^+^ 667.2, Found [M+H]^+^ 667.2. M.P: 78-80 ^o^C.

**Synthesis of PROTAC 13:**

***Tert-*butyl (6-aminohexyl)carbamate (S16):**

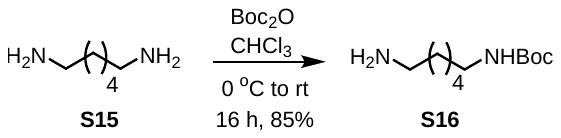

A solution of Boc_2_O (500 mg, 2.29 mmol, 1 equivalent) in CHCl_3_ (12 mL) was added dropwise to a solution of the diamine **S15** (2.1 g, 18.3 mmol, 8 equiv) in CHCl_3_ (18 mL) at 0 °C over a 30-min period. The reaction mixture was then stirred at room temperature for 16 h. The mixture was filtered, and the filtrate was concentrated under reduced pressure. The crude residue was dissolved in DCM (20 mL) and washed with brine (7 mL). The organic solution was dried over Na_2_SO_4,_ and the solution was filtered, concentrated and the residue was purified by silica gel column chromatography (1% Et_3_N in 0-5% MeOH/DCM) to give carbamate **S16** in 85% yield (419 mg) as a colorless oil.

^1^H NMR (400 MHz, CDCl_3_) *δ* 4.55 (s, 1H), 3.09 (q, *J* = 6.4 Hz, 2H), 2.66 (t, *J* = 6.8 Hz, 2H), 1.52 – 1.38 (m, 13H), 1.37-.28 (m, 4H).

**(*S*)-*N*-(6-Aminohexyl)-2-(4-(4-chlorophenyl)-2,3,9-trimethyl-6*H*-thieno[3,2-*f*][1,2,4]triazolo[4,3-*a*][1,4]diazepin-6-yl)acetamide (S18):**

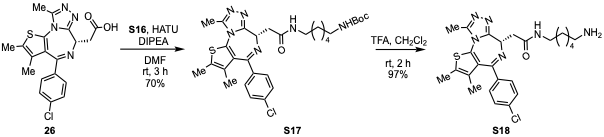

Acid **26** (100 mg, 0.25 mmol, 1.0 equivalents), DIPEA (174 µL, 4.0 equivalents) and HATU (190 mg, 2.0 equivalents) were added to a stirred solution of carbamate **S16** (65 mg, 0.3 mmol, 1.2 equivalents) in DMF (1.3 mL) at room temperature. The mixture was stirred for 3 h at room temperature. The mixture was then diluted with water (7 mL) and extracted with EtOAc (2 X 10 mL). The combined organic layers were washed with brine (5 mL), and the solution was dried over sodium sulfate. The solution was filtered, concentrated and the residue was purified by silica gel column chromatography (0-4% MeOH/DCM) to give compound **S17** in 70% (104 mg) as a yellowish solid.

^1^H NMR (400 MHz, CDCl_3_) *δ* 7.43 (d, *J* = 7.6 Hz, 2H), 7.35 (d, *J* = 7.6 Hz, 2H), 6.68 (brs, 1H), 4.67 (brs, 1H), 3.66 - 3.52 (m, 1H), 3.41 - 3.33 (m, 1H), 3.30 – 3.23 (m, 2H), 3.09 (brs, 2H), 2.70 (s, 3H), 2.41 (s, 3H), 1.68 (s, 3H), 1.58 – 1.49 (m, 4H), 1.44 (s, 9H), 1.34 – 1.28 (m, 4H). ESI-MS: Calcd for C_30_H_40_N_6_O_3_SCl [M+H]^+^ 599.2, Found [M+H]^+^ 599.1.

Boc-protected amine **S17** (48 mg, 0.07 mmol) was dissolved in DCM (1 mL). Then TFA (100 µL) (10:1 DCM:TFA) was added at room temperature. The reaction mixture was stirred for 2 hours at the same temperature. Afterward, the volatiles were evaporated, and the crude reaction mixture was dissolved in 10% MeOH in DCM (5 mL) and quenched with NaHCO_3_ (7 mL). The mixture was then extracted with 10% MeOH in DCM (2 × 10 mL), and the combined organic layers were dried over Na_2_SO_4_. The solution was filtered, concentrated under reduced pressure, and amine **S18** was obtained as a light-yellow solid in 97% (39 mg). This compound was used in subsequent reactions without further purification.

^1^H NMR (400 MHz, CDCl_3_) *δ* 7.40 (d, *J* = 8.4 Hz, 2H), 7.32 (d, *J* = 8.8 Hz, 2H), 6.88 (brs, 1H), 4.69 – 4.56 (m, 1H), 3.58 – 3.50 (m, 1H), 3.40 – 3.09 (m, 5H), 2.79 (brs, 2H), 2.66 (s, 3H), 2.40 (s, 3H), 1.67 (s, 3H), 1.60 – 1.49 (m, 4H), 1.45 – 1.29 (m, 4H). ESI-MS: Calcd for C_25_H_32_N_6_OSCl [M+H]^+^ 499.2, Found [M+H]^+^ 499.2.

**(*S*)-2-(4-Chlorophenoxy)-N-(6-(2-(4-(4-chlorophenyl)-2,3,9-trimethyl-6*H*-thieno[3,2-*f*][1,2,4]triazolo[4,3-*a*][1,4]diazepin-6-yl)acetamido)hexyl)-2-methylpropanamide (13):**

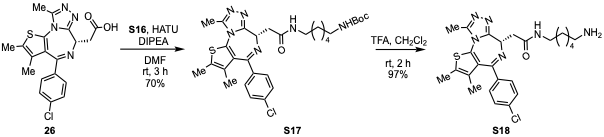

Amine **S18** (13 mg, 0.03 mmol, 1.0 equivalents) was dissolved in DMF (1 mL), followed by the addition of clofibric acid (**1**, 7 mg, 1.2 equivalents), DIPEA (18 µL, 4.0 equivalents), and HATU (20 mg, 2.0 equivalents) to the mixture at room temperature. The mixture was stirred for 2 hours at room temperature. The mixture was then diluted with water (5 mL) and extracted with EtOAc (2 X 10 mL). The combined organic layers were washed with brine (5 mL), and the solution was dried over sodium sulfate. The solution was filtered, concentrated, and the residue was purified by silica gel column chromatography (0-4% MeOH/DCM) to give compound **13** in 56% yield (10 mg) as a yellowish oil.

^1^H NMR (400 MHz, CDCl_3_) *δ* 7.45 (d, *J* = 8.0 Hz, 2H), 7.37 (d, *J* = 7.6 Hz, 2H), 7.22 (d, *J* = 8.8 Hz, 2H), 6.85 (d, *J* = 8.8 Hz, 2H), 6.78 (brs, 1H), 6.75 (brs, 1H), 4.70 (brs, 1H), 3.59 – 3.50 (m, 1H), 3.39 – 3.19 (m, 5H), 2.70 (s, 3H), 2.41 (s, 3H), 1.68 (s, 3H), 1.57 – 1.45 (m, 10H), 1.31 (brs, 4H). ^13^C NMR (101 MHz, CDCl_3_) *δ* 174.68, 170.69, 152.98, 131.58, 130.46, 129.38, 129.14, 128.58, 122.71, 81.95, 39.58, 39.30, 29.40, 29.29, 26.36, 26.31, 25.18, 14.54, 13.33. ESI-MS: Calcd for C_35_H_41_N_6_O_3_SCl_2_ [M+H]^+^ 695.2, Found [M+H]^+^ 695.2.

**Synthesis of PROTAC 14:**

***Tert-*butyl (8-aminooctyl)carbamate (S20):**

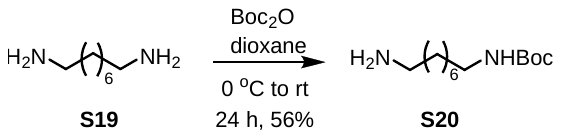

A solution of Boc_2_O (400 mg, 1.83 mmol, 1 equivalent) in THF (13 mL) was added dropwise to a solution of the diamine **S19** (2.0 g, 14.11 mmol, 8 equiv) in dioxane (20 mL) at 0 °C over a 30-min period. The reaction mixture was then stirred at room temperature for 24 h. The mixture was filtered, and the filtrate was concentrated in vacuo. The residue was resuspended in water (8 mL). The product was extracted with DCM (20 mL) and washed with brine (5 mL). The organic solution was dried over Na_2_SO_4,_ and the solution was filtered, concentrated and the residue was purified by silica gel column chromatography (1% NEt_3_ in 0-7% MeOH/DCM) to give carbamate **S20** in 56% yield (251 mg) as a light yellowish oil.

^1^H NMR (400 MHz, CDCl_3_) *δ* 4.52 (brs, 1H), 3.09 (q, *J* = 6.4 Hz, 2H), 2.68 (t, *J* = 6.8 Hz, 2H), 1.76 (brs, 2H), 1.43 (brs, 13H), 1.29 (brs, 8H).

**(*S*)-*N*-(8-Aminooctyl)-2-(4-(4-chlorophenyl)-2,3,9-trimethyl-6*H*-thieno[3,2-*f*][1,2,4]triazolo[4,3-*a*][1,4]diazepin-6-yl)acetamide (S22):**

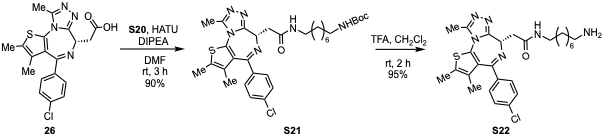

Acid **26** (100 mg, 0.25 mmol, 1.0 equivalents), DIPEA (174 µL, 4.0 equivalents) and HATU (190 mg, 2.0 equivalents) were added to a stirred solution of carbamate **S20** (65 mg, 0.3 mmol, 1.2 equivalents) in DMF (1.5 mL) at room temperature. The mixture was stirred for 3 h at room temperature. The mixture was then diluted with water (7 mL) and extracted with EtOAc (2 X 10 mL). The combined organic layers were washed with brine (5 mL), and the solution was dried over sodium sulfate. The solution was filtered, concentrated and the residue was purified by silica gel column chromatography (0-3% MeOH/DCM) to give compound **S21** in 90% (142 mg) as a dark-brown solid.

^1^H NMR (400 MHz, CDCl_3_) *δ* 7.42 (d, *J* = 8.3 Hz, 2H), 7.34 (d, *J* = 8.6 Hz, 2H), 6.58 (brs, 1H), 4.67 (t, *J* = 6.9 Hz, 1H), 4.55 (brs, 1H), 3.57 (dd, *J* = 14.0, 7.2 Hz, 1H), 3.42 – 3.18 (m, 3H), 3.09 (brs, 2H), 2.70 (s, 3H), 2.41 (s, 3H), 1.67 (s, 3H), 1.57 – 1.41 (m, 13H), 1.29 (brs, 8H). ESI-MS: Calcd for C_32_H_44_N_6_O_3_SCl [M+H]^+^ 627.3, Found [M+H]^+^ 627.3.

Boc-protected compound **S21** (137 mg, 0.22 mmol) was dissolved in DCM (1.8 mL).Then TFA (180 µL) (10:1 DCM:TFA) was added at room temperature. The reaction mixture was stirred for 2 hours at the same temperature. Afterward, the volatiles were evaporated, and the crude reaction mixture was dissolved in 10% MeOH in DCM (5 mL) and quenched with NaHCO_3_ (7 mL). The mixture was then extracted with 10% MeOH in DCM (2 × 10 mL), and the combined organic layers were dried over Na_2_SO_4_. The solution was filtered, concentrated under reduced pressure, and amine **S22** was obtained as a light-brown solid in 95% (110 mg). This compound was used in subsequent reactions without further purification.

^1^H NMR (400 MHz, CDCl_3_) *δ* 8.27 (brs, 2H), 7.39 (d, *J* = 8.4 Hz, 2H), 7.32 (dd, *J* = 8.4, 1.2 Hz, 2H), 7.21 (brs, 1H), 4.65 (t, *J* = 6.4 Hz, 1H), 3.52 (dd, *J* = 14.8, 8.0 Hz, 1H), 3.40 – 3.25 (m, 2H), 3.23 – 3.14 (m, 1H), 2.97 – 2.82 (m, 2H), 2.64 (s, 3H), 2.39 (s, 3H), 1.65 (s, 3H), 1.63 – 1.54 (m, 2H), 1.54 – 1.42 (m, 2H), 1.35 – 1.22 (m, 8H). ESI-MS: Calcd for C_27_H_36_N_6_OSCl [M+H]^+^ 527.2, Found [M+H]^+^ 527.2.

**(*S*)-2-(4-Chlorophenoxy)-*N*-(8-(2-(4-(4-chlorophenyl)-2,3,9-trimethyl-6*H*-thieno[3,2-*f*][1,2,4]triazolo[4,3-*a*][1,4]diazepin-6-yl)acetamido)octyl)-2-methylpropanamide (14):**

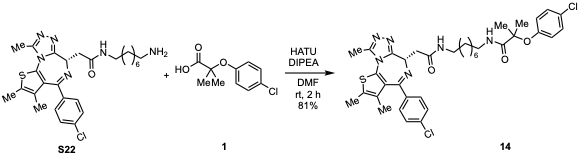

Amine **S22** (35 mg, 0.07 mmol, 1.0 equivalents) was dissolved in DMF (1 mL), followed by the addition of clofibric acid (**1**, 17 mg, 1.2 equivalents), DIPEA (46 µL, 4.0 equivalents), and HATU (51 mg, 2.0 equivalents) to the mixture at room temperature. The mixture was stirred for 2 hours at room temperature. The mixture was then diluted with water (5 mL) and extracted with EtOAc (2 X 10 mL). The combined organic layers were washed with brine (5 mL), and the solution was dried over sodium sulfate. The solution was filtered, concentrated, and the residue was purified by silica gel column chromatography (0-4% MeOH/DCM) to give compound **14** in 81% yield (39 mg) as a colorless semi-solid.

^1^H NMR (400 MHz, CDCl_3_) *δ* 7.40 (d, *J* = 8.4 Hz, 2H), 7.32 (d, *J* = 8.8 Hz, 2H), 7.25 – 7.19 (m, 2H), 6.88 – 6.81 (m, 2H), 6.66 (brs, 1H), 6.45 (brs, 1H), 4.61 (t, *J* = 6.8 Hz, 1H), 3.55 (dd, *J* = 14.4, 7.6 Hz, 1H), 3.38 – 3.17 (m, 5H), 2.66 (s, 3H), 2.40 (s, 3H), 1.67 (s, 3H), 1.55 – 1.43 (m, 4H), 1.49 (s, 6H), 1.26 (brs, 8H). ^13^C NMR (101 MHz, CDCl_3_) *δ* 174.37, 170.47, 164.08, 155.77, 153.06, 150.01, 137.00, 136.66, 132.30, 131.08, 131.04, 130.57, 129.98, 129.35, 128.87, 128.45, 122.49, 81.95, 54.66, 39.78, 39.62, 39.49, 29.59, 29.26, 26.93, 25.19, 14.52, 13.23, 11.97. ESI-MS: Calcd for C_37_H_45_N_6_O_3_SCl_2_ [M+H]^+^ 723.2, Found [M+H]^+^ 723.2.

**Synthesis of PROTAC 15:**

***Tert-*butyl (9-aminononyl)carbamate (S24):**

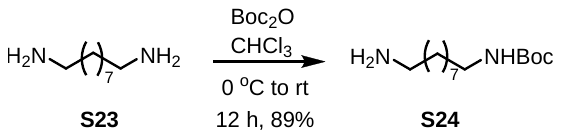

A solution of Boc_2_O (110 mg, 0.5 mmol, 1 equivalent) in CHCl_3_ (8 mL) was added dropwise to a solution of the diamine **23** (478 mg, 3.0 mmol, 6 equivalents) in CHCl_3_ (2 mL) at 0 °C over a 30 min period. The reaction mixture was then stirred at room temperature for 12 h. The mixture was filtered, and the filtrate was concentrated under reduced pressure. The crude residue was dissolved in DCM (15 mL) and washed with brine (5 mL). The organic solution was dried over Na_2_SO_4,_ and the solution was filtered, concentrated and the residue was purified by silica gel column chromatography (1% NEt_3_ in 0-5% MeOH/DCM) to give carbamate **S24** in 89% yield (116 mg) as a colorless semi-solid.

^1^H NMR (400 MHz, CDCl_3_) *δ* 4.51 (brs, 1H), 3.09 (q, *J* = 6.4 Hz, 2H), 2.69 (t, *J* = 7.0 Hz, 2H), 1.80 (brs, 2H), 1.43 (brs, 13H), 1.28 (brs, 10H).

***Tert-*butyl (9-(2-(4-chlorophenoxy)-2-methylpropanamido)nonyl)carbamate (S25):**

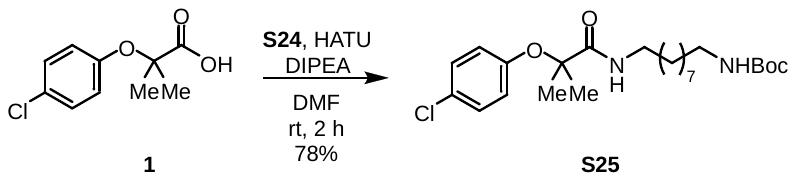

Acid **1** (70 mg, 0.33 mmol, 1.0 equivalent), DIPEA (227 µL, 4.0 equivalents) and HATU (186 mg, 1.5 equivalents) were added to a stirred solution of amine **S24** (101 mg, 0.39 mmol, 1.2 equivalents) in DMF (2 mL) at room temperature. The mixture was stirred for 2 h at room temperature. The mixture was then diluted with water (7 mL) and extracted with EtOAc (2 X 15 mL). The combined organic layers were washed with sat aq NH_4_Cl (5 mL), and the solution was dried over sodium sulfate. The solution was filtered, concentrated and the residue was purified by silica gel column chromatography (0-20% EA/Hex) to give compound **S25** in 78% (116 mg) yield as colorless oil.

^1^H NMR (400 MHz, CDCl_3_) *δ* 7.23 (d, *J* = 10.0 Hz, 2H), 6.84 (d, *J* = 10.4 Hz, 2H), 6.62 (brs, 1H), 4.50 (s, 1H), 3.28 (q, *J* = 6.8 Hz, 2H), 3.09 (q, *J* = 5.6 Hz, 2H), 1.53 – 1.42 (m, 19H), 1.26 (brs, 10H). ESI-MS: Calcd for C_24_H_40_ClN_2_O_4_ [M+H]^+^ 455.3, Found [M+H]^+^ 455.3.

**2-(4-Chlorophenoxy)-2-methyl-*N*-(9-((2,2,2-trifluoroacetyl)-l4-azaneyl)nonyl)propanamide (S26):**

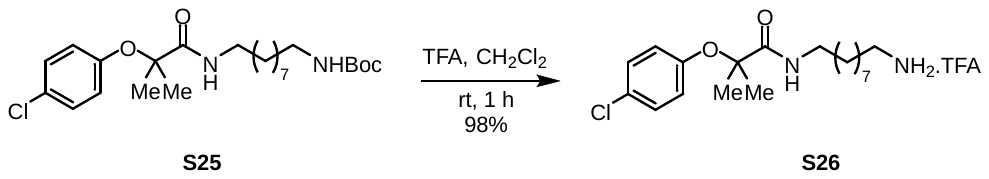

Compound **S25** (110 mg) was dissolved in DCM (1.5 mL). Then TFA (150 µL) (10:1 DCM:TFA) was added at room temperature. The reaction mixture was stirred for 1 hour at the same temperature. Afterward, the volatiles were then evaporated, and DCM (3 × 2 mL) was added to the residue. The mixture was re-evaporated three times. Finally, the residue was washed with hexanes and concentrated under reduced pressure, producing compound **S26** as a colorless oil in 98% yield (112 mg). This compound was used for next reaction without further purification.

^1^H NMR (400 MHz, CDCl_3_) *δ* 8.63 (brs, 3H), 7.28 – 7.22 (m, 2H), 6.99 (brs, 1H), 6.88 – 6.82 (m, 2H), 3.29 (q, *J* = 6.8 Hz, 2H), 3.03 (brs, 2H), 1.70 – 1.67 (m, 2H), 1.55 – 1.48 (m, 2H), 1.46 (s, 6H), 1.36 – 1.27 (s, 10H). ESI-MS: Calcd for C_19_H_32_ClN_2_O_2_ [M+H]^+^ 355.2, Found [M+H]^+^ 355.3.

**(*S*)-2-(4-Chlorophenoxy)-*N*-(9-(2-(4-(4-chlorophenyl)-2,3,9-trimethyl-6*H*-thieno[3,2-*f*][1,2,4]triazolo[4,3-*a*][1,4]diazepin-6-yl)acetamido)nonyl)-2-methylpropanamide (15):**

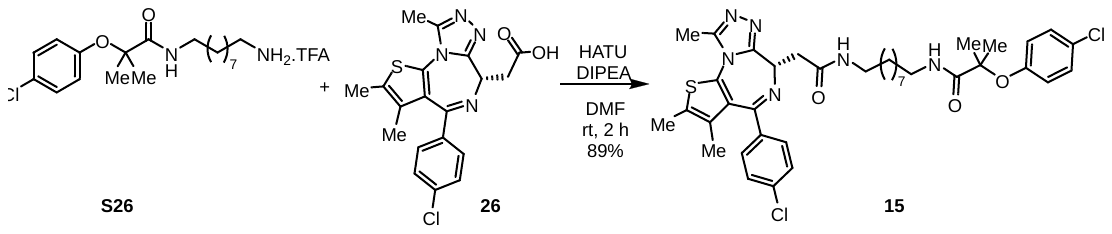

Acid **26** (25 mg, 0.06 mmol, 1.0 equivalents), DIPEA (43 µL, 4.0 equivalents) and HATU (36 mg, 1.5 equivalents) were added to a stirred solution of amine **S26** (35 mg, 0.07 mmol, 1.2 equivalents) in DMF (1 mL) at room temperature. The mixture was stirred for 2 h at room temperature. The mixture was then diluted with water (7 mL) and extracted with EtOAc (2 X 15 mL). The combined organic layers were washed with sat aq NH_4_Cl (5 mL), and the solution was dried over sodium sulfate. The solution was filtered, concentrated and the residue was purified by silica gel column chromatography (0-4% MeOH/DCM) to give compound **15** in 89% (41 mg) yield as colorless solid.

^1^H NMR (400 MHz, CDCl_3_) *δ* 7.39 (d, *J* = 8.4 Hz, 2H), 7.31 (d, *J* = 8.0 Hz, 2H), 7.21 (d, *J* = 8.8 Hz, 2H), 6.91 – 6.78 (m, 2H), 6.64 (brs, 1H), 6.50 (brs, 1H), 4.61 (t, *J* = 6.8 Hz, 1H), 3.53 (dd, *J* = 14.2, 7.4 Hz, 1H), 3.36 – 3.17 (m, 5H), 2.66 (s, 3H), 2.39 (s, 3H), 1.66 (s, 3H), 1.52 – 1.46 (m, 10H), 1.33 – 1.23 (m, 10H). ^13^C NMR (101 MHz, Chloroform-*d*) *δ* 174.34, 170.46, 164.02, 155.79, 153.03, 149.98, 136.94, 136.68, 132.26, 131.05, 130.99, 130.56, 129.96, 129.33, 128.83, 128.43, 122.48, 81.93, 54.62, 39.80, 39.54, 39.48, 29.60, 29.57, 29.49, 29.27, 26.99, 26.94, 25.17, 14.50, 13.21, 11.95. ESI-MS: Calcd for C_38_H_47_Cl_2_N_6_O_3_S [M+H]^+^ 737.3, Found [M+H]^+^ 737.2. M.P: 58-60 ^o^C.

**Synthesis of PROTAC 16:**

***Tert-*butyl (10-aminodecyl)carbamate (S28):**

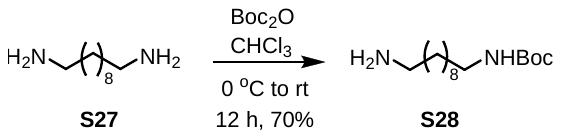

A solution of Boc_2_O (400 mg, 1.83 mmol, 1 equivalent) in CHCl_3_ (8 mL) was added dropwise to a solution of the diamine **S27** (1.6 g, 9.16 mmol, 5 equiv) in CHCl_3_ (30 mL) at 0 °C over a 30-min period. The reaction mixture was then stirred at room temperature for 12 h. The mixture was filtered, and the filtrate was concentrated under reduced pressure. The crude residue was dissolved in DCM (20 mL) and washed with brine (7 mL). The organic solution was dried over Na_2_SO_4,_ and the solution was filtered, concentrated and the residue was purified by silica gel column chromatography (1% Et_3_N in 0-5% MeOH/DCM) to give carbamate **S28** in 70% yield (349 mg) as a colorless semi-solid.

^1^H NMR (400 MHz, CDCl_3_) *δ* 4.50 (brs, 1H), 3.13 – 3.06 (m, 2H), 2.79 – 2.71 (m, 2H), 2.41 (brs, 2H), 1.51 – 1.41 (m, 13H), 1.27 (brs, 12H).

**(*S*)-*N*-(10-Aminodecyl)-2-(4-(4-chlorophenyl)-2,3,9-trimethyl-6*H*-thieno[3,2-*f*][1,2,4]triazolo[4,3-*a*][1,4]diazepin-6-yl)acetamide (30):**

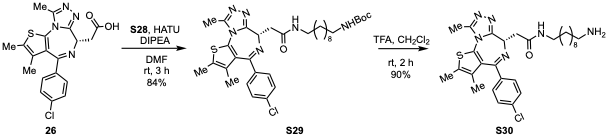

Acid **26** (100 mg, 0.25 mmol, 1.0 equivalents), DIPEA (174 µL, 4.0 equivalents) and HATU (190 mg, 2.0 equivalents) were added to a stirred solution of amine **S28** (82 mg, 0.3 mmol, 1.2 equivalents) in DMF (1.5 mL) at room temperature. The mixture was stirred for 3 h at room temperature. The mixture was then diluted with water (7 mL) and extracted with EtOAc (2 X 10 mL). The combined organic layers were washed with brine (5 mL), and the solution was dried over sodium sulfate. The solution was filtered, concentrated and the residue was purified by silica gel column chromatography (0-3% MeOH/DCM) to give compound **S29** in 84% (140 mg) as a yellow solid.

^1^H NMR (400 MHz, CDCl_3_) *δ* 7.40 (d, *J* = 8.4 Hz, 2H), 7.32 (d, *J* = 8.4 Hz, 2H), 6.47 (brs, 1H), 4.61 (t, *J* = 7.0 Hz, 1H), 4.52 (brs, 1H), 3.54 (dd, *J* = 14.0, 7.2 Hz, 1H), 3.36 – 3.15 (m, 3H), 3.13 – 3.04 (m, 2H), 2.66 (s, 3H), 2.40 (s, 3H), 1.66 (s, 3H), 1.57 – 1.37 (m, 13H), 1.26 (brs, 12H). ESI-MS: Calcd for C_34_H_48_N_6_O_3_SCl [M+H]^+^ 655.3, Found [M+H]^+^ 655.4.

Boc-protected amine **S29** (134 mg, 0.2 mmol) was dissolved in DCM (1.8 mL).Then TFA (180 µL) (10:1 DCM:TFA) was added at room temperature. The reaction mixture was stirred for 2 hours at the same temperature. Afterward, the volatiles were evaporated, and the crude reaction mixture was dissolved in 10% MeOH in DCM (5 mL) and quenched with NaHCO_3_ (7 mL). The mixture was then extracted with 10% MeOH in DCM (2 × 10 mL), and the combined organic layers were dried over Na_2_SO_4_. The solution was filtered, concentrated under reduced pressure, and amine **S30** was obtained as a light-yellowish solid in 90% (107 mg). This compound was used in subsequent reactions without further purification.

^1^H NMR (400 MHz, CDCl_3_) *δ* 7.39 (d, *J* = 8.4 Hz, 2H), 7.32 (d, *J* = 8.4 Hz, 2H), 7.04 (t, *J* = 5.6 Hz, 1H), 4.64 (t, *J* = 7.0 Hz, 1H), 3.48 (dd, *J* = 14.4, 7.2 Hz, 1H), 3.36 (dd, *J* = 14.4, 7.2 Hz, 1H), 3.31 – 3.20 (m, 2H), 2.84 (t, *J* = 7.5 Hz, 2H), 2.65 (s, 3H), 2.39 (s, 3H), 1.66 (s, 3H), 1.64 – 1.55 (m, 2H), 1.55 – 1.45 (m, 2H), 1.35 – 1.19 (m, 12H). ESI-MS: Calcd for C_29_H_40_N_6_OSCl [M+H]^+^ 555.2, Found [M+H]^+^ 555.3.

**(*S*)-2-(4-Chlorophenoxy)-*N*-(10-(2-(4-(4-chlorophenyl)-2,3,9-trimethyl-6*H*-thieno[3,2-*f*][1,2,4]triazolo[4,3-*a*][1,4]diazepin-6-yl)acetamido)decyl)-2-methylpropanamide (16):**

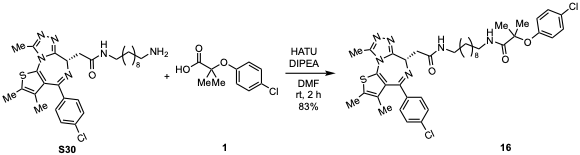

Amine **S30** (35 mg, 0.06 mmol, 1.0 equivalents) was dissolved in DMF (1 mL), followed by the addition of clofibric acid (**1**, 16 mg, 1.2 equivalents), DIPEA (44 µL, 4.0 equivalents), and HATU (48 mg, 2.0 equivalents) to the mixture at room temperature. The mixture was stirred for 2 hours at room temperature. The mixture was then diluted with sat. NH_4_Cl (5 mL) and extracted with EtOAc (2 X 10 mL). The solution was dried over sodium sulfate, concentrated, and the residue was purified by silica gel column chromatography (0-4% MeOH/DCM) to give compound **16** in 83% yield (39 mg) as a colorless semi-solid.

^1^H NMR (400 MHz, CDCl_3_) *δ* 7.40 (d, *J* = 8.4 Hz, 2H), 7.32 (d, *J* = 8.4 Hz, 2H), 7.23 – 7.19 (m, 2H), 6.89 – 6.79 (m, 2H), 6.64 (brs, 1H), 6.48 (brs, 1H), 4.62 (t, *J* = 6.4 Hz, 1H), 3.54 (dd, *J* = 14.0, 7.2 Hz, 1H), 3.39 – 3.16 (m, 5H), 2.66 (s, 3H), 2.40 (s, 3H), 1.66 (s, 3H), 1.55 – 1.45 (m, 10H), 1.35 – 1.17 (m, 12H). ^13^C NMR (101 MHz, CDCl_3_) *δ* 174.37, 170.51, 164.09, 155.76, 153.04, 150.03, 136.99, 136.66, 132.26, 131.09, 131.06, 130.59, 129.98, 129.35, 128.86, 128.46, 122.50, 81.95, 54.64, 39.84, 39.52, 29.63, 29.60, 29.55, 29.36, 29.35, 27.03, 26.99, 25.18, 14.51, 13.23, 11.95. ESI-MS: Calcd for C_39_H_49_N_6_O_3_SCl_2_ [M+H]^+^ 751.2, Found [M+H]^+^ 750.9.

**Synthesis of PROTAC 27:**

***N*-(4-Bromobenzyl)-2-(4-chlorophenoxy)-2-methylpropanamide (18):**

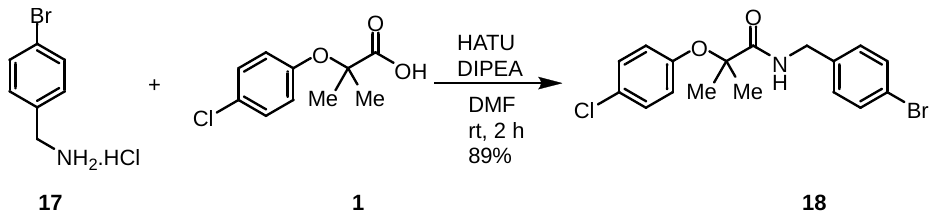

Acid **1** (200 mg, 0.93 mmol, 1.0 equivalent), DIPEA (649 µL, 4.0 equivalents) and HATU (532 mg, 1.5 equivalents) were added to a stirred solution of benzylamine **17** (250 mg, 1.12 mmol, 1.2 equivalents) in DMF (3 mL) at room temperature. The mixture was stirred for 2 h at room temperature. The mixture was then diluted with water (10 mL) and extracted with EtOAc (2 X 20 mL). The combined organic layers were washed with sat aq NH_4_Cl (10 mL), and the solution was dried over sodium sulfate. The solution was filtered, concentrated and the residue was purified by silica gel column chromatography (0-10% EtOAc in hexanes) to give compound **18** in 89% (319 mg) yield as colorless viscos oil.

^1^H NMR (400 MHz, CDCl_3_) *δ* 7.45 (dt, *J* = 8.8, 2.2 Hz, 2H), 7.21(dt, *J* = 8.8, 2.8 Hz, 2H), 7.13 (dt, *J* = 8.8, 2.0 Hz, 2H), 6.99 (brs, 1H), 6.81 (dt, *J* = 8.8, 2.8 Hz, 2H), 4.43 (d, *J* = 6.0 Hz, 2H), 1.51 (s, 6H).

**6-*tert*-Butyldimethylsilyloxy-1-hexyne (20)**.

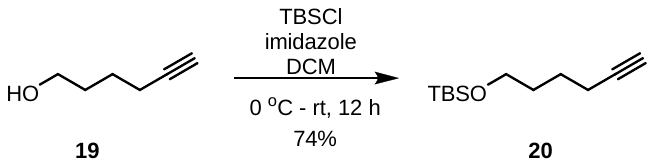

Hex-5-yn-1-ol (**19**, 500 mg, 5.1 mmol, 1.0 equiv) was dissolved in dichloromethane (8 mL) and the resulting solution was cooled to 0 °C. Imidazole (520 mg, 7.65 mmol, 1.5 equiv) was added, followed by tert-butyldimethylsilyl chloride (1.1 g, 7.65 mmol, 1.5 equiv) at the same temperature. The reaction mixture was allowed to warm to ambient temperature and was stirred for 12 h. The reaction was quenched by the addition of saturated aqueous NH₄Cl (15 mL) and the mixture was extracted with CH₂Cl₂ (2 × 15 mL). The combined organic layers were dried over anhydrous Na₂SO₄, filtered, and concentrated under reduced pressure to afford the TBS-protected compound 20 as a colorless oil in 74% yield (740 mg). This product was used directly for the next step without further purification.

***N*-(4-(6-((*Tert*-butyldimethylsilyl)oxy)hex-1-yn-1-yl)benzyl)-2-(4-chlorophenoxy)-2-methylpropanamide (21):**

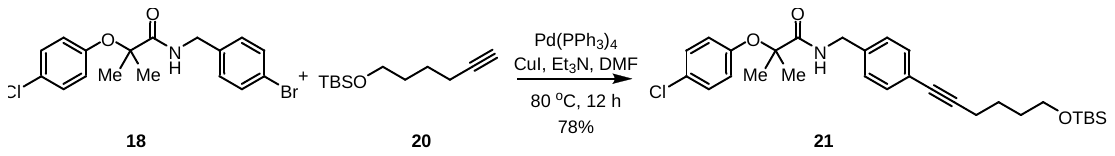

A round-bottom flask was charged with aryl bromide **18** (100 mg, 0.26 mmol, 1.0 equiv), alkyne **20** (72 mg, 1.3 equiv). Et₃N and DMF in a 1:1 ratio (3 mL total volume) was then added. The reaction mixture was purged with argon for approximately 15 minutes. Subsequently, Pd(PPh₃)₄ (2 mol%) and CuI (5 mol%) were added. The reaction mixture was then allowed to warm to 80 °C and stirred for 12 hours. Upon completion, the reaction was quenched with saturated ammonium chloride solution (10 mL). The mixture was extracted with ethyl acetate (2 × 15 mL). The combined organic layers were dried over anhydrous sodium sulfate, filtered, and concentrated under reduced pressure. The crude product was purified by silica gel column chromatography (0–15% ethyl acetate in hexanes) to afford compound **21** as an orange oil in 78% yield (106 mg).

^1^H NMR (400 MHz, CDCl_3_) *δ* 7.35 (d, *J* = 8.4 Hz, 2H), 7.21 (d, *J* = 10.0 Hz, 2H), 7.16 (d, *J* = 8.0 Hz, 2H), 6.96 (brs, 1H), 6.81 (d, *J* = 10.0 Hz, 2H), 4.46 (d, *J* = 5.6 Hz, 2H), 3.66 (t, *J* = 5.8 Hz, 2H), 2.43 (t, *J* = 6.6 Hz, 2H), 1.69 – 1.64 (m, 4H), 1.52 (s, 6H), 0.90 (s, 9H), 0.06 (s, 6H). ESI-MS: Calcd for C_29_H_41_ClNO_3_Si [M+H]^+^ 514.2, Found [M+H]^+^ 514.4.

**2-(4-Chlorophenoxy)-*N*-(4-(6-hydroxyhex-1-yn-1-yl)benzyl)-2-methylpropanamide (22):**

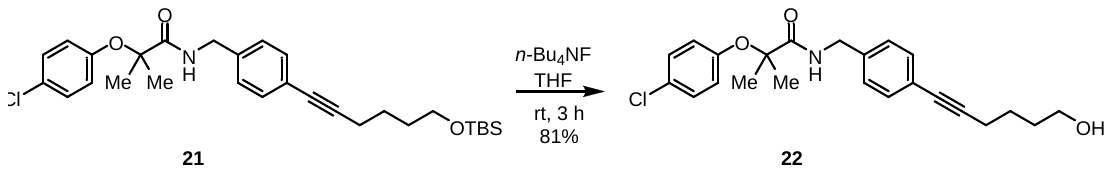

A TBS-protected compound **21** (104 mg, 0.2 mmol, 1.0 equiv) was dissolved in THF (4 mL), and a 1 M solution of tetrabutylammonium fluoride (TBAF, 607 μL, 3.0 equiv) was added at room temperature. The reaction mixture was stirred at the same temperature for 3 hours. Upon completion, the reaction mixture was diluted with ethyl acetate (20 mL) and washed sequentially with water (10 mL) and saturated sodium chloride solution (10 mL). The organic layer was separated, dried over anhydrous sodium sulfate, filtered, and concentrated under reduced pressure. The crude residue was purified by silica gel column chromatography (0–50% ethyl acetate in hexanes) to afford compound alcohol **22** as an orange oil in 81% yield (65 mg).

^1^H NMR (400 MHz, CDCl_3_) *δ* 7.34 (d, *J* = 8.4 Hz, 2H), 7.21 (d, *J* = 8.8 Hz, 2H), 7.16 (d, *J* = 8.0 Hz, 2H), 6.97 (brs, 1H), 6.81 (d, *J* = 8.4 Hz, 2H), 4.46 (d, *J* = 6.0 Hz, 2H), 3.71 (t, *J* = 6.2 Hz, 2H), 2.45 (t, *J* = 6.6 Hz, 2H), 1.80 – 1.65 (m, 4H), 1.51 (s, 6H). ESI-MS: Calcd for C_23_H_27_ClNO_3_ [M+H]^+^ 400.2, Found [M+H]^+^ 400.3.

**6-(4-((2-(4-Chlorophenoxy)-2-methylpropanamido)methyl)phenyl)hex-5-yn-1-yl 4-methylbenzenesulfonate (23):**

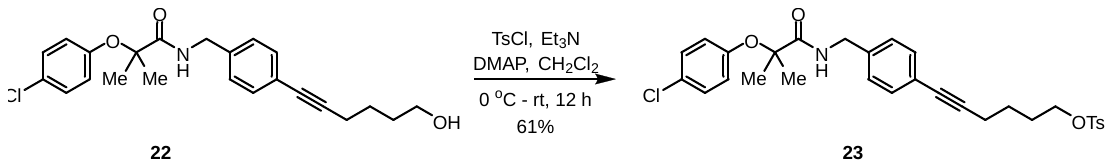

  Alcohol **22** (63 mg, 0.16 mmol, 1.0 equiv) was dissolved in CH₂Cl₂ (3 mL) under an argon atmosphere. Et₃N (44 μL, 2.0 equiv) and DMAP (4 mg, 0.2 equiv) were added to the reaction mixture at 0 °C. The mixture was stirred at the same temperature for 30 minutes. *p*-Toluenesulfonyl chloride (TsCl, 36 mg, 1.2 equiv) was then added in one portion at 0 °C. The reaction mixture was subsequently allowed to warm to room temperature and stir for 12 hours. Upon completion, the reaction was quenched with 20% aqueous citric acid solution (10 mL) and extracted with dichloromethane (20 mL). The organic layer was separated, dried over anhydrous sodium sulfate, filtered, and concentrated under reduced pressure. The crude product was purified by silica gel column chromatography (0–25% ethyl acetate in hexanes) to afford compound **23** as a colorless oil in 61% yield (53 mg).

^1^H NMR (400 MHz, CDCl_3_) *δ* 7.79 (d, *J* = 8.0 Hz, 2H), 7.35 – 7.29 (m, 4H), 7.21 (d, *J* = 10.0 Hz, 2H), 7.16 (d, *J* = 8.0 Hz, 1H), 6.97 (brs, 1H), 6.81 (d, *J* = 10.0 Hz, 2H), 4.46 (d, *J* = 6.0 Hz, 2H), 4.09 (t, *J* = 6.2 Hz, 2H), 2.44 (s, 3H), 2.38 (t, *J* = 6.8 Hz, 2H), 1.87 – 1.79 (m, 2H), 1.63 (quintet, *J* = 6.8 Hz, 2H), 1.51 (s, 6H). ESI-MS: Calcd for C_30_H_33_ClNO_5_S [M+H]^+^ 554.2, Found [M+H]^+^ 554.2.

***N*-(4-(6-Azidohex-1-yn-1-yl)benzyl)-2-(4-chlorophenoxy)-2-methylpropanamide (24):**

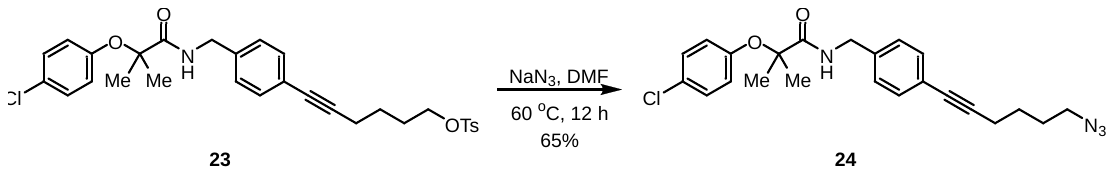

A round-bottom flask was charged with compound **23** (52 mg, 0.09 mmol, 1.0 equiv), NaN₃ (12 mg, 2.0 equiv), and a magnetic stir bar. The solids were dissolved in DMF (1.5 mL), and the reaction mixture was heated at 60 °C for 12 hours. After completion, the reaction mixture was cooled to 0 °C and diluted with water (10 mL). The resulting aqueous phase was extracted with diethyl ether (2 × 15 mL). The combined organic layers were washed with saturated brine solution (7 mL), dried over anhydrous sodium sulfate, filtered, and concentrated under reduced pressure. The crude residue was purified by silica gel column chromatography to afford compound **24** as a colorless oil in 65% yield (26 mg).

^1^H NMR (400 MHz, CDCl_3_) *δ* 7.34 (d, *J* = 8.0 Hz, 2H), 7.21 (d, *J* = 10.0 Hz, 2H), 7.17 (d, *J* = 8.4 Hz, 2H), 6.96 (brs, 1H), 6.81 (d, *J* = 10.0 Hz, 2H), 4.46 (d, *J* = 6.0 Hz, 2H), 3.34 (t, *J* = 6.7 Hz, 2H), 2.46 (t, *J* = 6.8 Hz, 2H), 1.82 – 1.73 (m, 2H), 1.71 – 1.65 (m, 2H), 1.52 (s, 6H).

***N*-(4-(6-Aminohex-1-yn-1-yl)benzyl)-2-(4-chlorophenoxy)-2-methylpropanamide (25):**

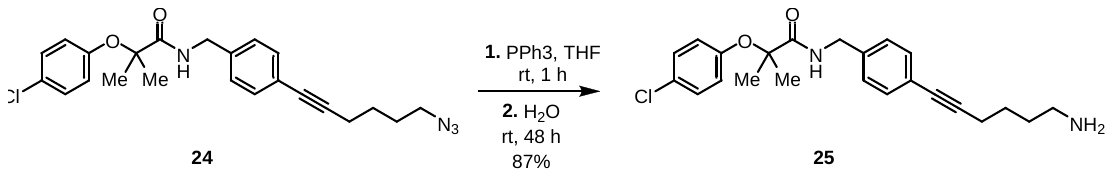

A round-bottom flask was charged with azide **24** (26 mg, 0.06 mmol, 1.0 equiv), anhydrous THF (1.0 mL), and a magnetic stir bar. A solution of PPh₃ (19 mg, 1.2 equiv) in THF (1.0 mL) was added to the reaction mixture. The resulting solution was stirred at room temperature for 1 hour, after which water (4 μL, 3.4 equiv) was added. The reaction mixture was stirred for an additional 48 hours. Upon completion, the solvent was evaporated under reduced pressure. The crude residue was purified by silica gel column chromatography to afford compound **25** as a colorless oil in 87% yield (21 mg).

^1^H NMR (400 MHz, CDCl_3_) *δ* 7.34 (d, *J* = 8.0 Hz, 2H), 7.20 (d, *J* = 10.0 Hz, 2H), 7.15 (d, *J* = 8.0 Hz, 2H), 6.98 (d, *J* = 5.4 Hz, 1H), 6.81 (dt, *J* = 10.0 Hz, 2H), 4.44 (d, *J* = 6.0 Hz, 1H), 2.88 (brs, 2H), 2.43 (t, *J* = 6.8 Hz, 2H), 1.76 – 1.71 (m, 2H), 1.70 – 1.63 (m, 2H), 1.51 (s, 6H). ESI-MS: Calcd for C_23_H_28_ClN_2_O_2_ [M+H]^+^ 399.2, Found [M+H]^+^ 399.2.

**(*S*)-2-(4-Chlorophenoxy)-*N*-(4-(6-(2-(4-(4-chlorophenyl)-2,3,9-trimethyl-6*H*-thieno[3,2-*f*][1,2,4]triazolo[4,3-*a*][1,4]diazepin-6-yl)acetamido)hex-1-yn-1-yl)benzyl)-2-methylpropanamide (27):**

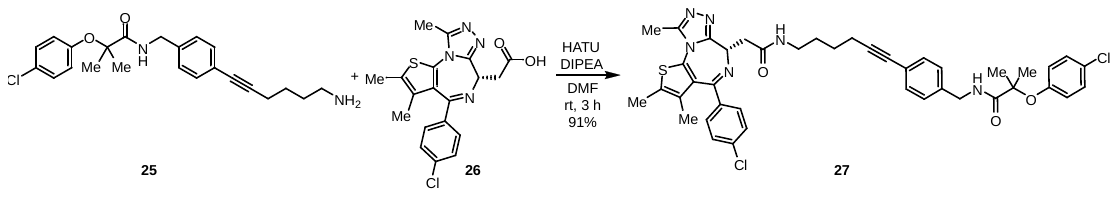

Acid **26** (17 mg, 0.04 mmol, 1.0 equivalent), DIPEA (30 µL, 4.0 equivalents) and HATU (24 mg, 1.5 equivalents) were added to a stirred solution of amine **25** (19 mg, 1.1 equivalents) in DMF (1.5 mL) at room temperature. The mixture was stirred for 3 h at room temperature. The mixture was then quenched with sat aq NH_4_Cl (7 mL) and extracted with EtOAc (2 X 15 mL). The combined organic layers were dried over sodium sulfate. The solution was filtered, concentrated and the residue was purified by silica gel column chromatography (0-3% MeOH in DCM) to give compound **27** in 91% (30 mg) yield as a colorless solid.

^1^H NMR (400 MHz, CDCl_3_) *δ* 7.40 (d, *J* = 8.0 Hz, 2H), 7.35 – 7.29 (m, 4H), 7.20 (d, *J* = 8.8 Hz, 2H), 7.15 (d, *J* = 8.0 Hz, 2H), 6.98 (brs, 1H), 6.81 (d, *J* = 8.8 Hz, 2H), 6.60 (brs, 1H), 4.63 (t, *J* = 6.2 Hz, 1H), 4.45 (d, *J* = 5.6 Hz, 2H), 3.55 (dd, *J* = 13.6, 6.8 Hz, 1H), 3.43 – 3.24 (m, 3H), 2.66 (s, 3H), 2.45 – 2.39 (m, 5H), 1.73 – 1.63 (m, 7H), 1.51 (s, 6H). ^13^C NMR (101 MHz, CDCl_3_) *δ* 174.47, 170.55, 164.19, 155.72, 152.81, 150.08, 137.61, 137.05, 136.55, 132.29, 132.02, 131.11, 130.56, 130.00, 129.38, 128.89, 128.72, 127.72, 123.30, 122.79, 90.15, 82.05, 80.75, 54.60, 43.27, 39.47, 39.31, 28.95, 26.15, 25.15, 19.26, 14.51, 13.22, 11.95. ESI-MS: Calcd for C_42_H_43_Cl_2_N_6_O_3_S [M+H]^+^ 781.2, Found [M+H]^+^ 781.3. M.P: 87 - 89 ^o^C.

**References**:

1. Eng, J. K.; Jahan, T. A.; Hoopmann, M. R. Comet: an open-source MS/MS sequence database search tool. *Proteomics* **2013,** *13* (1), 22-4, 10.1002/pmic.201200439.

2. Wilmarth, P. A.; Riviere, M. A.; David, L. L. Techniques for accurate protein identification in shotgun proteomic studies of human, mouse, bovine, and chicken lenses. *J. Ocul. Biol. Dis. Infor.* **2009,** *2* (4), 223-234, 10.1007/s12177-009-9042-6.

3. Robinson, M. D.; McCarthy, D. J.; Smyth, G. K. edgeR: a Bioconductor package for differential expression analysis of digital gene expression data. *Bioinformatics* **2010,** *26* (1), 139-40, 10.1093/bioinformatics/btp616.
